# Non-antibiotic pharmaceuticals can enhance the spread of antibiotic resistance via conjugation

**DOI:** 10.1101/724500

**Authors:** Yue Wang, Ji Lu, Shuai Zhang, Jie Li, Likai Mao, Zhiguo Yuan, Philip L. Bond, Jianhua Guo

## Abstract

Antibiotic resistance is a global threat for public health. It is widely acknowledged that antibiotics at sub-inhibitory concentrations are important in disseminating antibiotic resistance via horizontal gene transfer. While there is high use of non-antibiotic human-targeted pharmaceuticals in our societies, the potential contribution of these on the spread of antibiotic resistance has been overlooked so far. Here, we report that commonly consumed non-antibiotic pharmaceuticals, including nonsteroidal anti-inflammatories (ibuprofen, naproxen, diclofenac), a lipid-lowering drug (gemfibrozil), and a *β*-blocker (propanolol), at clinically and environmentally relevant concentrations, significantly accelerated the conjugation of plasmid-borne antibiotic resistance genes. We looked at the response to these drugs by the bacteria involved in the gene transfer through various analyses that included monitoring reactive oxygen species (ROS) and cell membrane permeability by flow cytometry, cell arrangement, and whole-genome RNA and protein sequencing. We found the enhanced conjugation correlated well with increased production of ROS and cell membrane permeability. We also detected closer cell-to-cell contact and upregulated conjugal genes. Additionally, these non-antibiotic pharmaceuticals caused the bacteria to have responses similar to those detected when exposed to antibiotics, such as inducing the SOS response, and enhancing efflux pumps. The findings advance our understanding of the bacterial transfer of antibiotic resistance genes, and importantly emphasize concerns of non-antibiotic human-targeted pharmaceuticals for enhancing the spread of antibiotic resistance.

## Introduction

Increasing levels of antibiotic resistance occurring in bacteria is seen to be a major threat for human health, which is put forward by World Health Organization. Currently, this is causing at least 700 000 deaths worldwide annually ^1^. The acquisition of antibiotic resistance can mainly occur through a mutation in bacterial DNA or by obtaining antibiotic resistance genes (ARGs) through horizontal gene transfer (HGT) ^2, 3^. HGT consists of three different pathways: conjugation, transformation and transduction. Among them, conjugation is a main mechanism for disseminating antibiotic resistance ^4^. During conjugation, the exchange of genetic material between the donor and recipient bacteria occurs by direct cell-to-cell contact and by a connecting pilus ^5^. Typically, the exchange is mediated by mobile genetic elements, such as a conjugative plasmid.

It is commonly acknowledged that the emergence and spread of antibiotic resistance is largely due to misuse and overuse of antibiotics in clinical, veterinary, and agricultural settings ^6^. Exposure of microorganisms to antibiotics that are below the minimal inhibitory concentration (MIC) can promote HGT ^7, 8^. For example, antibiotics aminoglycoside and fluoroquinolone were shown to induce genetic transformability in pathogen *Streptococcus pneumonia*e ^7^. Although the consumption of non-antibiotic pharmaceuticals occupy approximately 95% of the drug market ^9, 10^, the role of non-antibiotic pharmaceuticals in the emergence and spread of antibiotic resistance has received relatively little attention. Recently, Maier et al. ^11^ screened more than 1 000 marketed drugs against 40 representative gut bacterial strains, and reported that more than 200 non-antibiotic pharmaceuticals could exhibit antibiotic-like effects on the bacteria. They found these non-antibiotic pharmaceuticals contributed to the emergence of antibiotic resistance through mutation or increased expression of efflux pump genes ^11^. However, they did not investigate if these non-antibiotic pharmaceuticals can facilitate HGT ^11^. If non-antibiotic pharmaceuticals have effects on the spread of antibiotic resistance, there may be features and properties of the non-antibiotic pharmaceuticals, or shared mechanisms, that promote the horizontal transfer of ARGs.

In this study, we investigated the potential of different types of commonly consumed non-antibiotic human-targeted pharmaceuticals for promoting conjugative transfer of plasmid borne ARGs. We also explored the underlying mechanisms contributing to the HGT. The tested pharmaceuticals were the nonsteroidal anti-inflammatory drugs (NSAIDs) (ibuprofen, naproxen, diclofenac), a lipid-lowering drug (gemfibrozil), a *β*-blocker (propanolol), and a contrast medium (iopromide). The use of these drugs covers a wide range of clinical settings that includes pain/fever-relief, inflammatory-treatment, lipid control, heart disease, and diagnostic medicine. All these pharmaceuticals are on the World Health Organization’s List of Essential Medicines and are highly consumed. For example, there are 30 million worldwide-users of NSAIDs daily, and over 100 million annual consumptions in the USA alone ^12^. Such drugs are presented in the human gut or plasma at high concentrations. Levels of diclofenac and ibuprofen can be at 2.2 mg/L and 11.4 mg/L in plasma, respectively ^13, 14^, while gemfibrozil is reported to occur at 17.8 mg/L in plasma ^15^. In addition, after human administration, a large portion of these drugs (e.g., up to 90%) is excreted unchanged in the urine and ultimately ends up in wastewater ^16, 17^. Thus, these pharmaceuticals are also recognized as emerging contaminants and are ubiquitously detected in various environments, including wastewater, surface water, groundwater, and even drinking water, ranging in concentrations from nanograms to milligram per litre ^18, 19^.

Here we showed that five non-antibiotic pharmaceuticals, ibuprofen, naproxen, gemfibrozil, diclofenac, and propranolol, significantly facilitated conjugative transfer of plasmid-borne ARGs across bacterial genera; while iopromide did not. The phenotypic (culture-based plating and fluorescence-based flow cytometry) and genotypic (plasmid electrophoresis, whole-genome RNA sequencing and proteomic analysis) data provided collective evidence of the underlying mechanisms for the increased HGT. The five non-antibiotic pharmaceuticals, with antibiotic-like effects towards bacteria, induced over-production of reactive oxygen species (ROS), increased cell membrane permeability, facilitated cell-to-cell contact, and modulated conjugal genes (including upregulation of pilin generation). We propose these responses to the drugs boosted the frequency of HGT of the ARGs. The findings increase our insight of the spread of antibiotic resistance, and suggest the antibiotic-like potential of non-antibiotic pharmaceuticals should not be overlooked for drug development.

## Results

### Non-antibiotic pharmaceuticals significantly accelerate conjugative transfer of ARGs

To evaluate the effects of six non-antibiotic pharmaceuticals on conjugation, we used *E. coli* LE392 with the conjugative RP4 plasmid harbouring multiple resistance genes against tetracycline, kanamycin, and ampicillin as the donor. *Pseudomonas putida* KT2440, with high tolerance towards chloramphenicol, was the recipient ^20^. During the conjugation process the cells were exposed to sub-inhibitory non-antibiotic pharmaceuticals (MICs towards pharmaceuticals are shown in Table S1), at concentrations from 0.0001 to 50 mg/L (both clinical and environmentally relevant concentrations were included) to test if they could increase the transfer of ARGs. After the cross genera mating, transconjugants were distinguished and enumerated on plates containing four antibiotics (tetracycline, kanamycin, ampicillin and chloramphenicol). The transfer events in the different treatment groups were enumerated as the absolute number of transconjugants, and normalized as the transfer frequency, which was calculated as the number of transconjugants divided by the number of recipients (Fig. 1a).

**Fig 1.**
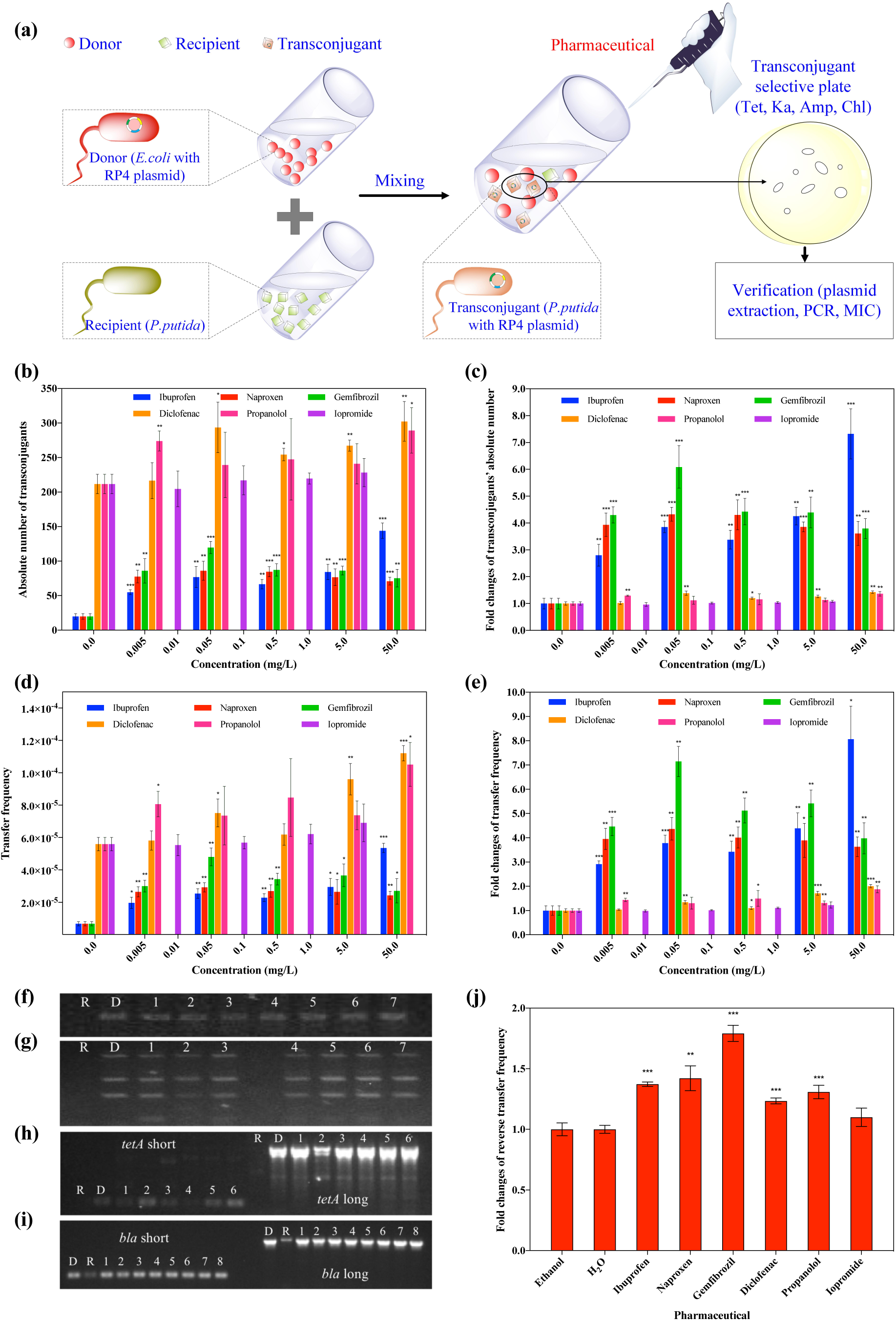
Effects of non-antibiotic pharmaceuticals on the conjugative transfer of ARGs. (a) Schematic experimental design of the conjugation. (b) Absolute number of transconjugants under the exposure of non-antibiotic pharmaceuticals. (c) Fold changes of transconjugants’ absolute number. (d) Transfer frequency under the exposure of non-antibiotic pharmaceuticals. (e) Fold changes of transfer frequency under the exposure of non-antibiotic pharmaceuticals. (f) Electrophoresis of RP4 plasmid (lanes R, D, and 1-7, refer to plasmids extracted from recipient, donor, and transconjugants of different pharmaceutical-dosed groups). (g) Electrophoresis of RP4 plasmid detection using specific primers (lanes R, D, and 1-7, refer to plasmids extracted from recipient, donor, and transconjugants of different pharmaceutical-dosed groups). (h) Electrophoresis of plasmid PCR products for *tetA* gene (lanes R, D, and 1-6, refer to plasmids extracted from recipient, donor, and transconjugants of different pharmaceutical-dosed groups). (i) Electrophoresis of plasmid PCR products for *bla* gene (lanes R, D, and 1-8, refer to plasmids extracted from recipient, donor, and transconjugants of different pharmaceutical-dosed groups). (j) Fold changes of reverse transfer frequency under the exposure of non-antibiotic pharmaceuticals (0.5 mg/L for ibuprofen, naproxen, gemfibrozil, diclofenac, propanolol, and 1.0 mg/L for iopromide). Significant differences between non-antibiotic-dosed samples and the control were analyzed by independent-sample *t* test, \**P* < 0.05, \*\**P* < 0.01, and \*\*\**P* < 0.001.

In term of both the number of transconjugants and the transfer frequency, we found that with the addition of non-antibiotic pharmaceuticals, ibuprofen, naproxen and gemfibrozil, at all five concentrations used here (from 0.005 to 50 mg/L), the conjugative transfer increased significantly (*P* < 0.05). For diclofenac and propanolol, only the higher concentrations (5 or 50 mg/L) enhanced conjugative transfer. In contrast, none of the applied iopromide concentrations increased the conjugation (Fig. 1b-d). Using ibuprofen as a specific example, the absolute number of transconjugants increased from 20±4 to 144±11 when increasing its dosage from 0 mg/L to 50 mg/L. Regarding the transfer frequency, the spontaneous frequency was low, this being 5.6×10^−5^±4.2×10^−6^ and 6.8×10^−6^±1.3×10^−6^ for MilliQ water and ethanol, respectively. According to fold changes of conjugative transfer frequency, except for iopromide, the five non-antibiotic pharmaceuticals, at concentrations as low as 0.05 mg/L, increased transfer frequencies significantly (*P* < 0.05) (Fig. 1e). The fold change could be as high as 8 times when exposed to 50 mg/L ibuprofen for 8 h. Even lower concentration (0.005 mg/L) of ibuprofen, naproxen and gemfibrozil also showed significant enhancement in the conjugative frequency.

To verify the successful transfer of the RP4 plasmid, gel electrophoresis showed that the plasmids in transconjugants were the same as that in the donor while no plasmid was seen in the recipient (Fig. 1f), and the specific primers used generated three bands also showed the same result (Fig. 1g). PCR of *tetA* and *bla* genes (both short and long primers applied) also indicated the plasmids from transconjugants harboured these same genes as that in donor (Fig. 1h and Fig. 1i). MICs of the transconjugants towards the four antibiotics, tetracycline, kanamycin, ampicillin, and chloramphenicol, were the same as those of the donor and recipient bacterium (Table S2).

We found also that the RP4 plasmid was able to transfer from the transconjugant to the recipient bacterium *E. coli* MG1655 ^21^. In addition, when exposing the reverse mating system to ibuprofen, naproxen, gemfibrozil, diclofenac, and propanolol at 0.5 mg/L, the fold changes of transfer frequency were significantly increased (*P* < 0.05) (Fig. 1j).

Collectively, it can be concluded that the non-antibiotic pharmaceuticals (excepting for iopromide) significantly increased intergenera conjugative transfer of the multiresistance genes (*P* < 0.05). In addition, the generated transconjugant is able to transfer the plasmid and could become a new source of ARGs.

### ROS play an important role in the enhanced conjugative transfer

ROS are natural byproducts of metabolism in bacteria. However, under environmental stress, ROS production may increase dramatically, and this may enhance conjugative transfer ^20, 22^. We hypothesized the increased conjugation frequency is due to raised ROS levels. Consequently, in conjugation experiments as described above, the fluorescence-measured ROS production was seen to increase significantly in both the donor and recipient under exposure of the five non-antibiotic pharmaceuticals (except for iopromide) (*P* < 0.05) (Fig. S1). Noticeably, the solvent ethanol (with 1% final volume ratio) did not increase ROS levels significantly compared to the solvent MilliQ water. By comparing to the corresponding control group, the fold changes of ROS levels in the donor bacteria increased from 2-fold, to up 15-fold at exposure of 50 mg/L propanolol (Fig. 2a). The fold changes of ROS generation in the recipient was relatively lower than those in the donor, in which the highest change was 3-fold at the exposure of 50 mg/L ibuprofen (Fig. 2b). Moreover, the effects of diclofenac and propanolol on ROS generation in the donor were concentration-dependent (*r*=0.84, *P* < 0.05 and *r*=0.86, *P* < 0.01 for diclofenac and propanolol, respectively), higher ROS levels were detected with increasing concentrations of pharmaceuticals. In contrast the effects of ibuprofen, naproxen and gemfibrozil exhibited a concentration-independent effect on ROS (*P* > 0.05). It should be noted that the increase of ROS generation was due to the dosage of pharmaceuticals, based on the fact that ethanol did not increase the ROS generation.

**Fig 2.**
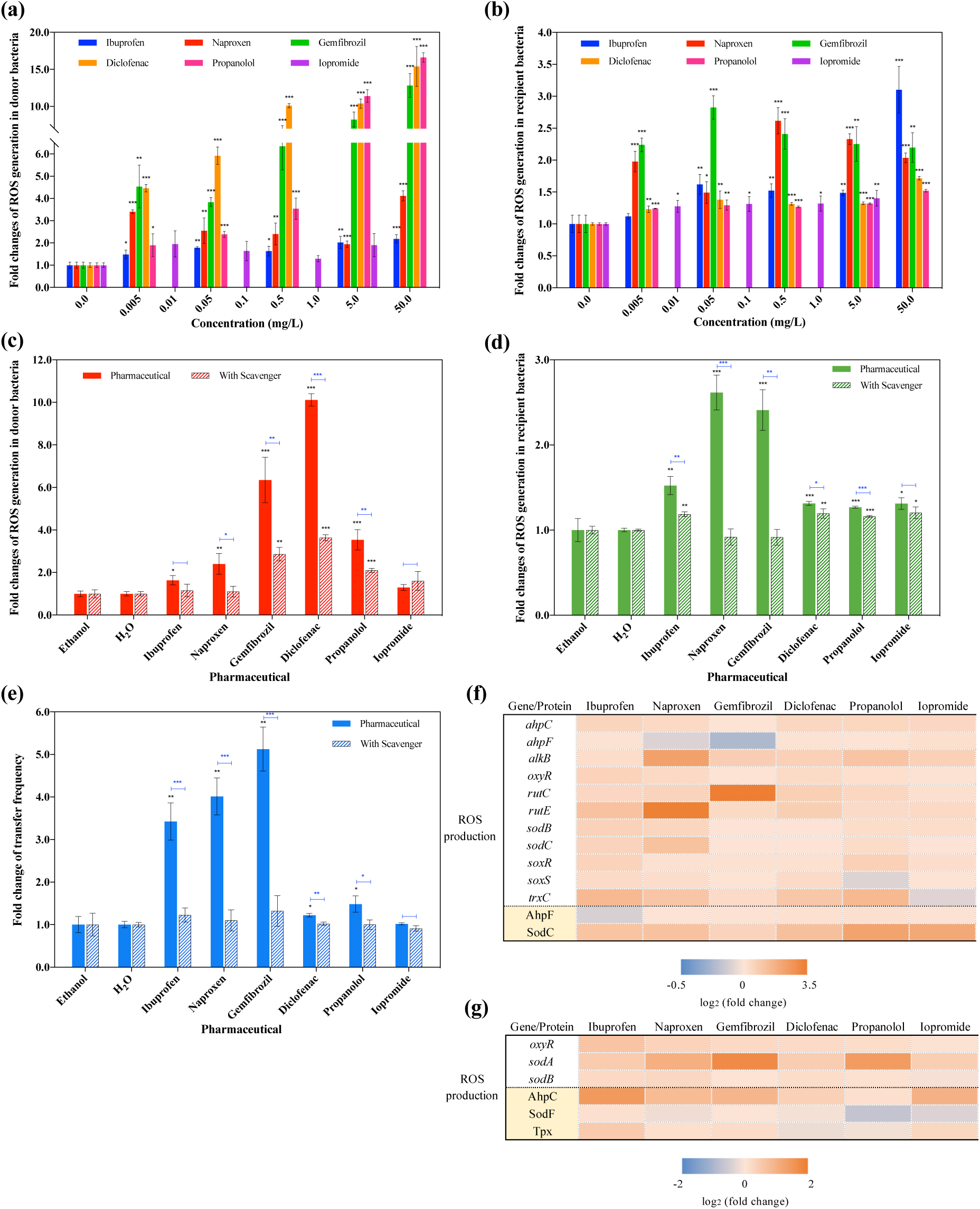
Effects of non-antibiotic pharmaceuticals on ROS in the donor (*E. coli* K-12 LE392) and recipient (*P. putida* KT2440) bacteria. (a) Fold changes of ROS generation in donor bacteria. (b) Fold changes of ROS generation in recipient bacteria. (c) Fold changes of ROS generation in donor bacteria with the addition of ROS scavenger thiourea. (d) Fold changes of ROS generation in recipient bacteria with the addition of ROS scavenger thiourea. (e) Fold changes of conjugative transfer frequency with the addition of ROS scavenger thiourea. (f) Fold changes of expression of core genes and proteins related to ROS production in donor bacteria. (g) Fold changes of expression of core genes and proteins related to ROS production in recipient bacteria. Significant differences between non-antibiotic-dosed samples and the control were analyzed by independent-sample *t* test, \**P* < 0.05, \*\**P* < 0.01, and \*\*\**P* < 0.001. For (c)-(g), figures shown are 0.5 mg/L for ibuprofen, naproxen, gemfibrozil, diclofenac, propanolol, and 1.0 mg/L for iopromide.

We found that an ROS scavenger (thiourea) could eliminate the over-production of ROS, caused by non-antibiotic pharmaceuticals, in both donor and recipient bacteria (*P* < 0.05) (Fig. 2c, 2d, Fig. S1). With the exception that 0.5 mg/L of diclofenac and propanolol could still significantly increase ROS generation in both the donor and recipient (*P* < 0.05, Fig. 2c, 2d). In that case there may be some other ROS produced that are not eliminated by thiourea. Nonetheless, we were able to experimentally reverse the effects of ROS on the conjugation process, by adding thiourea during the mating period. As illustrated in Fig. 2e and Fig. S1, the conjugative transfer frequency declined significantly for all the pharmaceuticals (*P* < 0.05) in the presence of the scavenger. For example, with 0.5 mg/L gemfibrozil and naproxen, the fold change of transfer frequency decreased from 5-fold and 4-fold to only 1.3-fold and 1.1-fold, respectively, when the scavenger was added. No significant increase was observed in the transfer frequency between the controls (no drug) and the scavenger-dosed drug groups (Fig. 2e), indicating that the ROS are playing an important role in the pharmaceutical enhanced conjugation process.

In the conjugation experiments we compared expression levels of RNA and protein between the non-antibiotic pharmaceuticals dosed groups and the control groups (no drugs applied) of the donor and recipient bacteria. This was conducted to further understand the effects of these pharmaceuticals on conjugation. It was seen that the pharmaceuticals enhanced ROS production-related proteins and genes significantly in both donor and recipient (Fig. 2f, Fig. 2g, Tables S3-S6). For the donor bacterium, these pharmaceuticals enhanced expression of redox-sensing genes, *oxyR* and *soxR*, which are the regulators of genes for defending oxidative stress ^23, 24^ (Fig. 2f). Proteins responsible for alkyl hydroperoxide reductase (AhpF) and superoxide dismutase (SodC) activities increased significantly with the dosage of pharmaceuticals (*q* < 0.01). For example, expression of SodC was enhanced 4.7-fold when exposed to 0.5 mg/L propanolol. Correspondingly, genes coding for hydroperoxide reductase (*ahpC* and *ahpF*), oxidative demethylase (*alkB*), superoxide dismutase (*sodB* and *sodC*) and superoxide response (*soxS*) increased under the exposure of pharmaceuticals by 1.1- to 4.8-fold. These genes are involved in the bacterial response to high-level oxidative stress ^25^. Noticeably, iopromide of 1.0 mg/L had the least effect on the ROS-related gene expression levels in the donor bacterium, which is in agreement with lower levels of ROS generation detected for that exposure (Fig. 2a). For the recipient bacterium, these non-antibiotic pharmaceuticals increased protein abundances of alkyl hydroperoxide reductase (AhpF) and hydroperoxide peroxidase (Tpx), but only ibuprofen and gemfibrozil enhanced the expression of superoxide dismutase protein (SodF) (Fig. 2g). Additionally, the expression of the redox-sensing gene (*oxyR*) and the superoxide dismutase regulators (*sodA* and *sodB*), were significantly enhanced under the exposure of all pharmaceuticals.

### Cell membrane variations link to increased conjugation

If the cell membranes become more permeable, it will be easier for plasmid to transfer from donor to recipient bacteria during the conjugative process ^26^. We speculate that non-antibiotic pharmaceuticals might increase conjugative transfer by affecting cell membrane. Thus, we tested the cell membrane permeability by flow cytometry in the bacteria in the presence and absence of the pharmaceuticals. For the donor bacteria, naproxen, gemfibrozil, diclofenac, and propanolol at the low concentration of 0.005 mg/L were seen to increase the cell membrane permeability significantly (*P* < 0.05) (Fig. 3a and Fig. S2). Ibuprofen at concentrations higher than 0.05 mg/L significantly increased the membrane permeability, while iopromide had no effect (Fig. 3a). The impact of ibuprofen on the donor bacteria’s cell membrane permeability was concentration-dependent (*r*=0.98, *P* < 0.01), such that the membrane permeability increased with the increase of ibuprofen, and a 2.5-fold change was detected at 50.0 mg/L. In contrast, for the other non-antibiotic pharmaceuticals, the membrane permeability changes were not seen to be concentration-dependent (*P* > 0.05). The results matched well with the conjugative transfer changes detected, where the frequency was more enhanced with increasing ibuprofen concentrations (Fig. 1a and 1b). For the recipient bacteria, all the chosen concentrations of ibuprofen, naproxen, gemfibrozil, diclofenac, and propanolol enhanced the membrane permeability significantly (*P* < 0.05) (Fig. 3b and Fig. S2). These increases in cell membrane permeability are likely contributing to the increased conjugation detected for these non-antibiotic pharmaceuticals.

**Fig 3.**
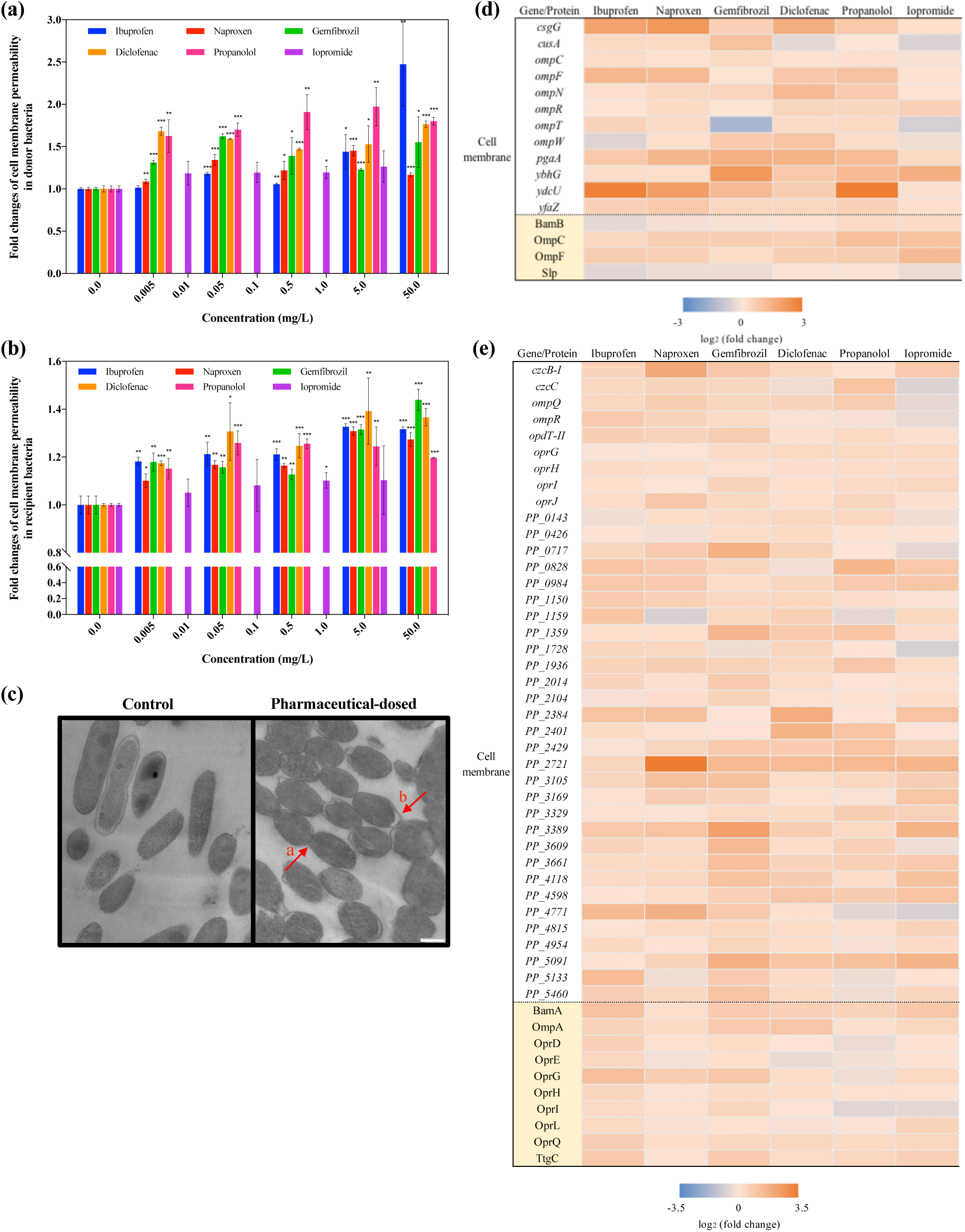
Effects of non-antibiotic pharmaceuticals on cell membranes in the donor (*E. coli* K-12 LE392) and recipient (*P. putida* KT2440) bacteria. (a) Fold changes of cell membrane permeability in donor bacteria. (b) Fold changes of cell membrane permeability in recipient bacteria. (c) TEM images of donor and recipient bacteria under the exposure of pharmaceuticals. Cells remained separate and intact in the control group; while cells became closer (arrow a) and membranes were partially damaged (arrow b) with pharmaceutical dosage. (d) Fold changes of expression of core genes and proteins related to cell membranes in donor bacteria. (e) Fold changes of expression of core genes and proteins related to cell membranes in recipient bacteria. Significant differences between non-antibiotic-dosed samples and the control were analyzed by independent-sample *t* test, \**P* < 0.05, \*\**P* < 0.01, and \*\*\**P* < 0.001. For (d)-(e), figures shown are 0.5 mg/L for ibuprofen, naproxen, gemfibrozil, diclofenac, propanolol, and 1.0 mg/L for iopromide.

We examined the effect of the pharmaceuticals on the cell morphology and arrangement during the conjugation periods by transmission electron microscopy (TEM). During exposure to the pharmaceuticals (excepting for iopromide) the cells became more compact and closer (Arrow a in Fig. 3c), and cell membranes were partially damaged (Arrow b in Fig. 3c). In contrast, for iopromide, the cells remained separate and intact (Fig. S3). During the conjugation process direct donor and recipient cell contact is a necessity for the plasmid transfer ^27^. Thus, the closer cell contact and membrane damage detected here agrees with the changes in membrane permeability and the correspondingly higher levels of gene transfer detected in the presence of the pharmaceuticals. This provides further explanation for the enhanced conjugative transfer detected for ibuprofen, naproxen, gemfibrozil, diclofenac and propranolol, and is in agreement with the lack of effect by iopromide.

Moreover, the variations of cell membranes induced by non-antibiotic pharmaceuticals were supported by the analyses at both RNA and protein levels. Core genes and proteins related to cell membrane structure and function showed significant changes under the exposure of the non-antibiotic pharmaceuticals (Tables S7-S10). Regulator proteins, which alter the levels of outer membrane channels and membrane permeability ^28, 29^, increased significantly after exposure to the non-antibiotic pharmaceuticals (*q* < 0.01). For example, OmpC and OmpF in the donor bacteria, and OmpA, OprH, OprL and OprQ in the recipient bacteria, showed significant enhancement of abundance in all of the five pharmaceutical-dosed groups (Fig. 3d and 3e). The increase was as high as 2.4-fold. The correspondingly relevant genes also showed significantly increased expression. This included *ompC*, *ompF*, *ompN*, *ompR* in the donor bacteria, and *oprG*, *oprH*, *oprI*, *oprJ* in the recipient bacteria. Noticeably, the expression of the genes *ompC*, *ompF*, *ompN* in the donor bacteria were not changed for iopromide, while the other five pharmaceuticals caused up to 2.5-fold change. A decrease in expression of *ompQ* and *ompR* genes was detected in the recipient bacteria after dosing iopromide, whereas ibuprofen, naproxen, gemfibrozil, diclofenac and propanolol caused their increased expression from 1.3-1.8 folds. These variations also partially explain the different effects of pharmaceuticals on the conjugation process. In addition, putative genes which code for outer membrane proteins in donor bacteria ^30^, also increased significantly due to the effects of non-antibiotic pharmaceuticals. For example, the genes *csgG*, *cusA*, *pgaA*, *ybhG*, *ydcU*, *yfaZ* had increased expression by up to 8 folds (with iopromide exposure had the least increase effect), and these may also be contributing to the increased cell membrane permeability.

### Other key factors regulating conjugative process

Genes on the conjugative plasmid are also key factors in regulating conjugation, which involves coordinated processes of replication, partitioning and conjugation ^31^. In particular, the global regulator *korB* alters operon expression of the IncP-*α* RP4 plasmid. Under the exposure of these non-antibiotic pharmaceuticals the expression of *korB* was repressed by 1.1- to 1.7-fold decrease (Fig. 4a), thus, leading to the enhanced expression of genes for the mating-pair apparatus, replication and conjugative regulators. For example, ibuprofen at 0.5 mg/L caused enhanced expression of the conjugative transfer transcriptional regulator, *traG* and *trbD* by up to 2.2- and 1.7-fold, respectively; caused up-regulation of the mating-pair apparatus, including *trbA*, *trbK*, *trfA2*, by up to 235-fold; and it increased expression of the replication regulator, where a 2-fold change in *traC1* was detected. Similar changes were seen when the RP4 plasmid was exposed to naproxen, gemfibrozil, diclofenac, and propanolol. Noticeably, iopromide had the least effect on *korB* expression, with only a 1.1-fold decrease, thus, having lower effect on other core genes in RP4 plasmid. For example, expression of *trfA2* gene, which is responsible for mating pair formation and replication in the RP4 plasmid ^32, 33^, showed a 15-fold decrease under the effect of iopromide. However, the expression of the gene was enhanced by 56 to 271 folds when exposed to the other five pharmaceuticals (Table S11). As for the other factors influencing the transfer frequency, this also partially explains why iopromide was less effective in promoting conjugal process.

**Fig 4.**
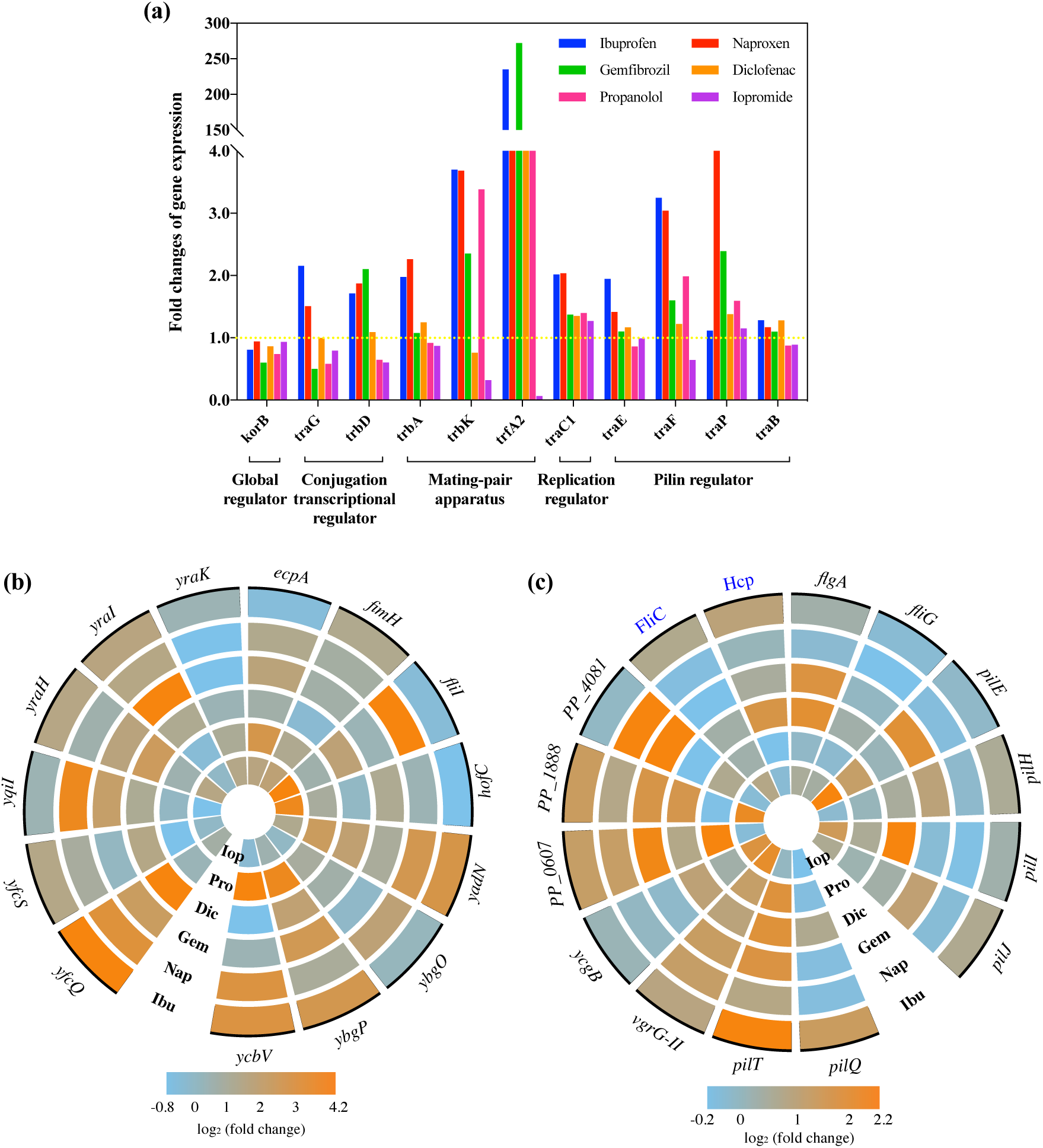
Effects of non-antibiotic pharmaceuticals on fimbriae gene expression in the donor (*E. coli* K-12 LE392), recipient (*P. putida* KT2440) bacteria, and core gene expression in conjugative plasmid (IncP-α RP4 plasmid). (a) Fold changes of expression of core genes in RP4 plasmid. (b) Fold changes of expression of core genes related to fimbriae in donor bacteria. (c) Fold changes of expression of core genes and proteins related to fimbriae in recipient bacteria. Ibu, Nap, Gem, Dic, Pro, and Iop refer to 0.5 mg/L ibuprofen, 0.5 mg/L naproxen, 0.5 mg/L gemfibrozil, 0.5 mg/L diclofenac, 0.5 mg/L propanolol, and 1.0 mg/L iopromide, respectively.

During the conjugation process the plasmid is transferred through a pilin bridge, and the pilin-related genes in RP4 plasmid include *traB*, *traE*, *traF*, and *traP* ^34^. Under the exposure of ibuprofen, naproxen, gemfibrozil and diclofenac, all these four genes were up-regulated by 1.1- to 15-fold enhancement compared to the control group. For propanolol, increased expression of *traF* and *traP* to 2-fold was detected, but decreased expression levels of *traB* and *traE* to 1.2-fold occurred. Significant increases of pilin gene expression were not detected for iopromide exposure, although we observed decreased expression of *traB*, *traE* and *traF*.

Another contributing factor to conjugation is the direct cell-to-cell contact ^27^, to which fimbriae are important for bacterial cell adhesion. Fimbriae generation and functions are regulated within the regulator operons *fim*, *pil*, *yad*, *ybg*, *ycb*, *yfc*, *yra*, *ycg* ^35–37^. In this study, genes and proteins related to fimbriae adhesion were up-regulated significantly under the exposure of the five non-antibiotic pharmaceuticals, excluding the effect of iopromide (Tables S12-S14). For example, in the donor bacteria the gene expression was enhanced by as high as 17.8-fold under the effect of 0.5 mg/L gemfibrozil (Fig. 4b). While in the recipient bacteria, the highest increase was to 0.5 mg/L naproxen, with a 4.3-fold increase (Fig. 4c). In comparison, iopromide exposure repressed expression of most of the fimbriae-related genes in the donor bacteria by 1.2 to 1.8 folds.

### Antibiotic-like features caused by non-antibiotic pharmaceuticals

Antibiotics at sub-inhibitory concentrations are known to promote horizontal dissemination of antibiotic resistance, which is associated with the SOS response of bacteria ^6, 8^. In this study, we found the non-antibiotic pharmaceuticals also had significant effects on SOS response in both donor and recipient bacteria (Fig. 5, Tables S15-S18). Altered gene expression during pharmaceutical exposure was detected for the key regulators of *lexA*, *umu*, *yeb* in the donor, and *sox* in the recipient, with a total of seven genes being affected ^38^. Under the exposure of ibuprofen, naproxen, and propanolol, all of these core genes responsible for SOS response had increased expression by 1.1 to 4.2 folds. While gemfibrozil and diclofenac caused enhanced expression of six of the seven genes, with the largest change being 5.4-fold. In contrast, iopromide caused increased expression of three of the seven genes, which were *umuD* in the donor (1.1-fold), and *soxD* (1.5-fold) and *soxR* in the recipient (4.0-fold); and caused decreased expression of the other four genes by 1.1- to 1.5-fold. Thus, the SOS response could also contribute to the non-antibiotic pharmaceutical-enhanced conjugation, and help explain the differences detected under the exposure of different pharmaceuticals.

**Fig 5.**
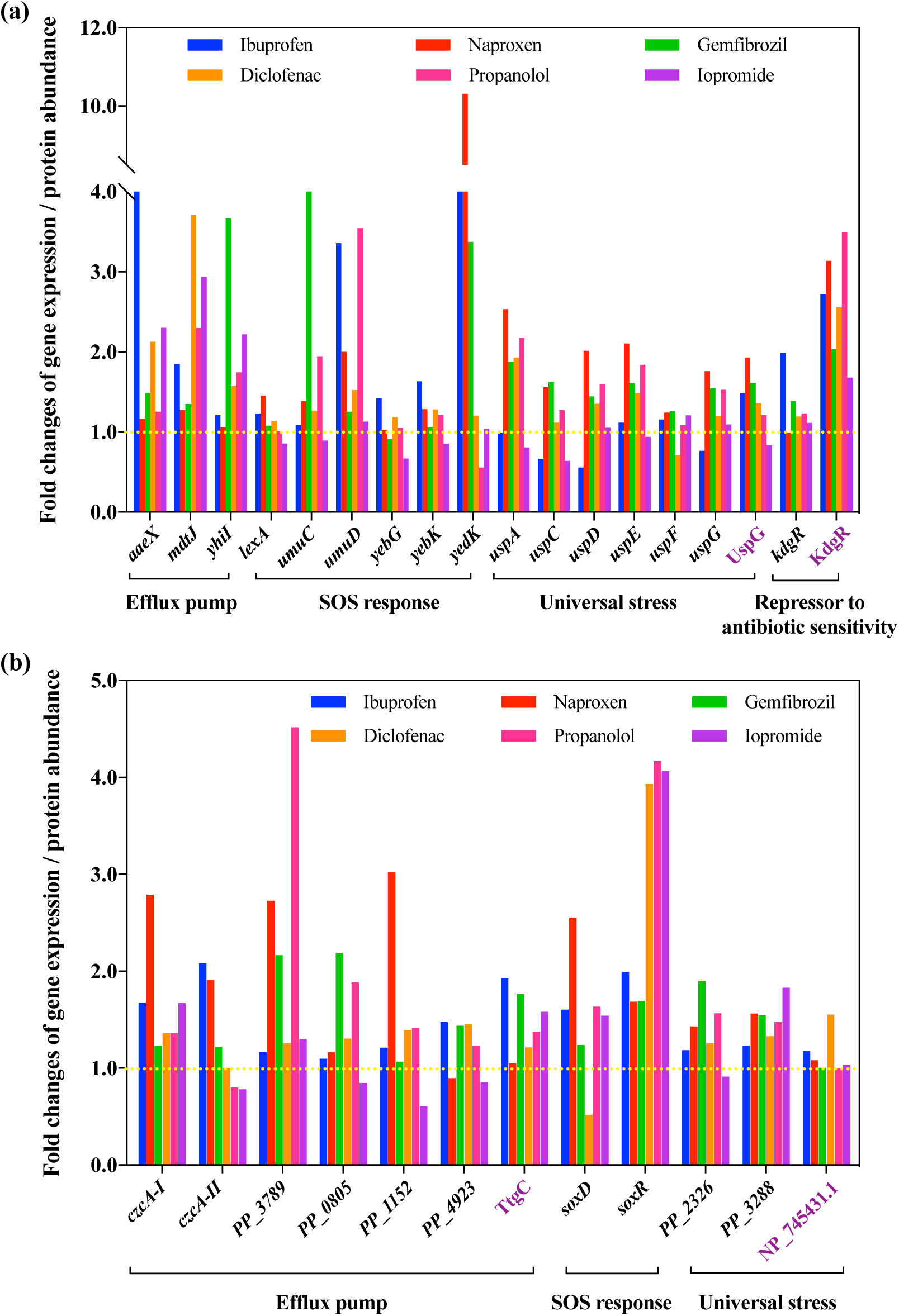
Non-antibiotic pharmaceuticals showed antibiotic-like features on donor (*E. coli* K-12 LE392) and recipient (*P. putida* KT2440) bacteria. (a) Fold changes of expression of core genes and proteins in donor bacteria. (b) Fold changes of expression of core genes and proteins in recipient bacteria. Figures shown are 0.5 mg/L for ibuprofen, naproxen, gemfibrozil, diclofenac, propanolol, and 1.0 mg/L for iopromide. Genes are shown in black, while proteins are shown in purple.

In addition to the SOS response, these non-antibiotic pharmaceuticals also had influence on other effects that antibiotics may cause on both the donor and recipient, this included the enhanced expression of efflux pumps, increased levels of universal stress, and even elevated levels of repressor genes which regulate antibiotic-sensitivity. Core operons of these effects are *mdt*, *usp*, *kdg* in donor, and *czc*, *ttg* in the recipient bacteria ^39, 40^. Despite some fluctuations, these five non-antibiotic pharmaceuticals caused increased expression of the relevant genes; while exposure to 1.0 mg/L iopromide showed the least effects on changed gene expression (Tables S19-S20).

## Discussion

Pharmaceuticals are being consumed at alarmingly increased levels in recent years. The global pharmaceuticals market was worth $935 billion in 2017, and will reach $1170 billion in 2021, with a 5.8% yearly growth ^9, 10^. Among the highly-consumed pharmaceuticals, antibiotic consumption is only $43 billion, which occupies a 4.6% portion of the market. The dominant portion of the market is non-antibiotic pharmaceuticals ^9, 10^. It is well studied that antibiotics at sub-inhibitory concentrations can facilitate the spread of antibiotic resistance ^41–45^. However, the contribution of non-antibiotic human-targeted pharmaceuticals on the spread of antibiotic resistance have been severely overlooked. In this study, we demonstrated that the exposure of bacteria to five commonly consumed non-antibiotic human-targeted pharmaceuticals (ibuprofen, naproxen, gemfibrozil, diclofenac, and propanolol) caused increased dissemination of antibiotic resistance via conjugative transfer. In contrast, the diagnostic drug, iopromide, did not result in increased gene transfer. The changes of absolute number of transconjugants and the transfer frequency both increased significantly under the exposure of ibuprofen, naproxen, gemfibrozil with the concentrations as low as 0.005 mg/L, or in the presence of diclofenac, propanolol with concentrations higher than 0.05 mg/L. Noticeably, we further confirmed successful transfer of the RP4 plasmid by PCR of plasmid genes, testing the antibiotic MIC of the transconjugants, and conducting reverse transfer from the transconjugants. These findings enabled ruling out any co-selective effects or mutagenesis, and coincided with the phenotypic results. Compared with the conjugation effects caused by sub-inhibitory antibiotics, the fold changes were comparable, or lower, for example, sub-inhibitory tetracycline in drinking water resulted in a 10-fold increasement for the transfer of the conjugative element from *Enterococcus faecalis* to *Listeria monocytogenes*^46^. However, considering the consumption is relatively high, the effects caused by non-antibiotic pharmaceuticals cannot be ignored. Moreover, this is the first time to report that these five commonly consumed non-antibiotic pharmaceuticals (ibuprofen, naproxen, gemfibrozil, diclofenac, and propanolol) can enhance the spread of antibiotic resistance under both clinical- and environmentally-relevant concentrations.

Additionally, in this study we explored the underlying mechanisms relating to the increased gene transfer by culturing- and fluorescence-based methods ^21, 47^, as well as by advanced molecular techniques (Fig. 6). The higher levels of ROS triggered by these non-antibiotic pharmaceuticals is likely a major influence on the increased gene transfer. Under the exposure of ibuprofen, naproxen, gemfibrozil, diclofenac, and propanolol, intracellular ROS production was increased significantly (*P* < 0.05). Both RNA and protein levels indicated significant increased expression of oxidative regulators, *oxyR* and *soxR*, and this coincided with the over-expression of antioxidant genes, including superoxide dismutase *sod* and hydroperoxide reductase *ahp* (*P* < 0.05) ^23, 24^. After adding the ROS scavenger, these non-antibiotic pharmaceuticals did not cause enhanced intracellular ROS generation for both the donor and recipient. Consequently, the enhanced conjugative transfer frequency was eliminated by addition of the ROS scavenger. In addition, iopromide did not promote the conjugative transfer, likely because it did not cause ROS stress in the donor and recipient cells.

**Fig 6.**
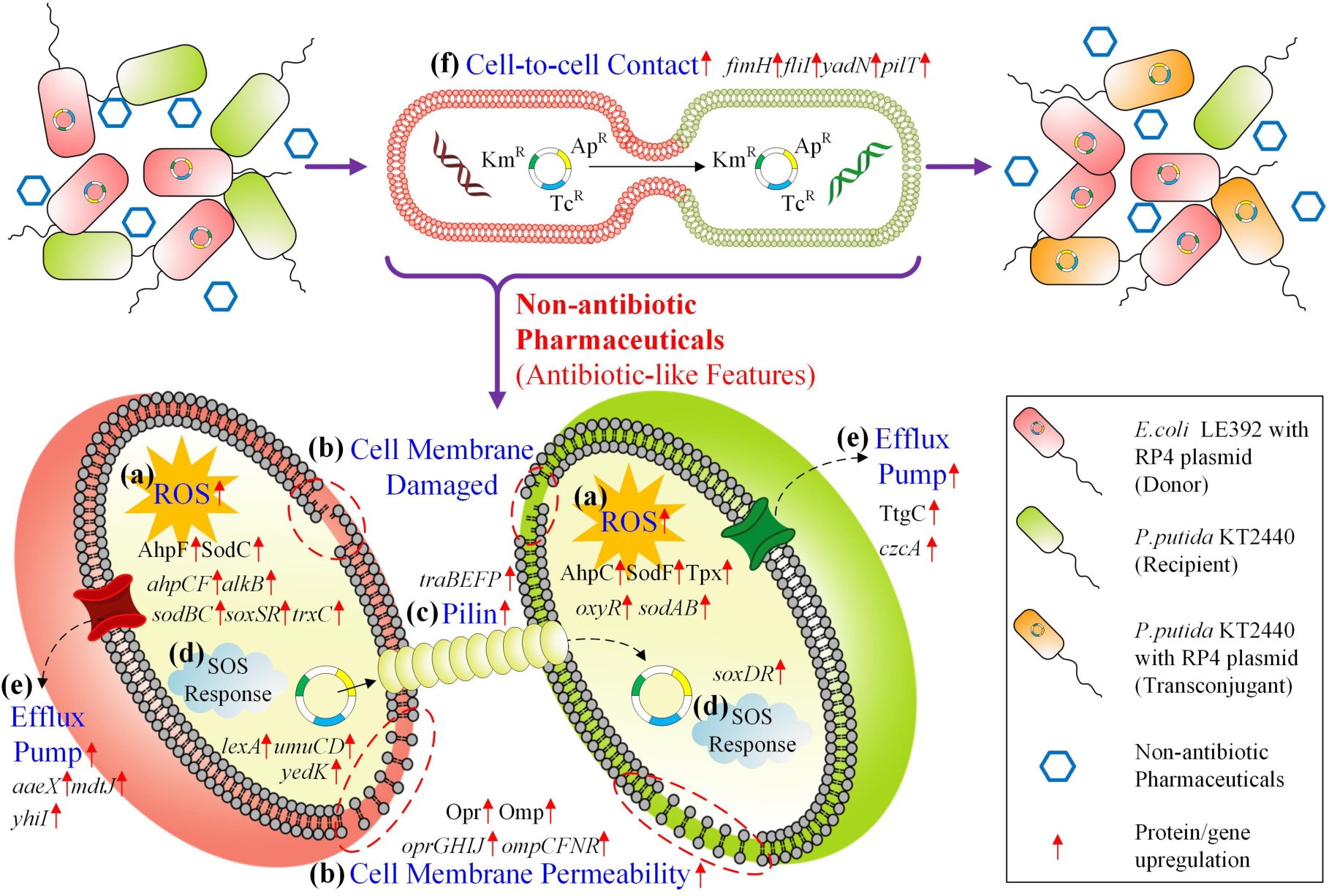
The overall mechanisms of non-antibiotic human-targeted pharmaceuticals causing increased conjugative transfer of plasmid-borne ARGs. (a) Non-antibiotic pharmaceuticals enhance ROS production in both donor and recipient bacteria. (b) Non-antibiotic pharmaceuticals induce cell membrane variations, including increasing cell membrane permeability and causing cell membrane damage in both donor and recipient bacteria. (c) Non-antibiotic pharmaceuticals promote pilin generation in donor bacterial strain. (d) SOS response in both donor and recipient bacteria was triggered under the exposure of non-antibiotic pharmaceuticals. (e) Efflux pump in both donor and recipient bacteria was activated in the presence of non-antibiotic pharmaceuticals. (f) Non-antibiotic pharmaceuticals facilitate cell-to-cell contact.

We also found that the condition of the cell membrane is an important factor for facilitating conjugation by detecting changes in cell membrane permeability and observing cell-to-cell contact. Elevated cell membrane permeability was detected in both the donor and recipient cells under the exposure of ibuprofen, naproxen, gemfibrozil, diclofenac, and propanolol. On the contrary, iopromide did not cause similar effects. Transcriptional and protein expression levels also supported these findings. These exposures caused increased levels of outer membrane regulon proteins Omp and Opr, together with the corresponding up-regulated *omp* and *opr* genes, while iopromide exposure caused lower levels of change. The outer membrane of Gram-negative bacteria is considered to be a semi-permeable barrier, where increased permeability could enable increased entry of plasmids ^48^. It is also reported that the transient membrane permeability has evolutionary implications and can facilitate horizontal gene transfer ^49^. Additionally, direct cell-to-cell contact is required for transfer of plasmids from donor to recipient via pilin bridge ^27^. In this study, TEM indicated that ibuprofen, naproxen, gemfibrozil, diclofenac, and propanolol could promote cell contact, while iopromide did not. We also found that the enhanced levels of fimbriae-related proteins and genes may play a role. Fimbriae is reported to increase cell adhesion and promote the formation of biofilms ^50^. In this study, iopromide had the least effect on fimbriae-gene/protein regulations compared to the other five pharmaceuticals. Therefore, the variations of cell membrane integrity, permeability and cell-to-cell contact is likely contributing to the enhanced conjugation (Fig. 6).

For the RP4 plasmid important plasmid borne factors for the conjugative process are the DNA-transfer replication (Dtr) and the mating pair formation (Mpf) systems ^51^. The Dtr system is essential for plasmid replication and the Mpf system is responsible for the generation of pilin ^52^. Upon exposure to the non-antibiotic pharmaceuticals significant variations of both the Dtr and Mpf systems were detected. For Dtr, the *traC* gene was up-regulated in the presence of non-antibiotic pharmaceuticals. For the Mpf system, the genes *trbK*, *trfA* (mating-pair apparatus), and *traF*, *traP* (pilin regulator), had increased levels of expression under the exposure of ibuprofen, naproxen, gemfibrozil, diclofenac, and propanolol; while decreased levels were observed when exposing to iopromide. Thus, we propose that variation of the RP4 plasmid gene expression, caused by the pharmaceutical exposure, is contributing to the enhanced conjugative transfer (Fig. 6).

Interestingly, we also found these non-antibiotic pharmaceuticals caused antibiotic-like bacterial responses. Here we detected the increased expression of genes and proteins involved in the SOS response (*lexA*, *umuC*, *umuD* and *soxR*), universal stress (Usp), efflux pump (*aaeX*, *mdtJ*, *yhiI* and *czcA*), and antibiotic-sensitivity (KdgR). Other *in vivo* studies show that some pharmaceuticals can cause stress on cells. For example in humans, ibuprofen could enhance oxidative stress in plasma during extreme exercise ^53^ and induce prolonged stress in a rat model ^54^. Naproxen can induce oxidative stress and genotoxicity in male Wistar rats ^55^. Diclofenac is also demonstrated to possess a broad antimicrobial activity *in vitro* ^56^. It is also reported that human-targeted non-antibiotic drugs boost antibiotic-like side effects on the gut microbiome ^11^. Here they detected that bacterial mutant strains lacking TolC, which is responsible for efflux of antibiotics, became more sensitive to antibiotics and human-targeted non-antibiotic drugs.

We aimed to determine some key features of these non-antibiotic pharmaceuticals contributing to the stimulatory effects on the conjugative process. Our results indicate that non-antibiotic pharmaceuticals that cause increased intracellular ROS generation will likely cause increased gene transfer by conjugation. In addition to these five non-antibiotic pharmaceuticals reported in this study, we previously also reported that carbamazepine could facilitate the conjugative transfer due to enhanced ROS production ^20^. Previous studies also documented that these non-antibiotic pharmaceuticals cause negative effects on the health status in animals and humans due to oxidative stress. For example, NSAID-pharmaceuticals (e.g., ibuprofen, naproxen, and diclofenac) have been reported to induce cardiotoxicity by a ROS-dependent mechanism, and were further verified with the addition of antioxidants ^57–59^. Therefore, these studies on animals or humans support the increase in ROS in the presence of these non-antibiotic pharmaceuticals. In addition to these non-antibiotic pharmaceuticals, biocides (e.g., triclosan) and heavy metals were also demonstrated to increase ROS generation levels, impose stress-response on bacteria, thus enhancing the uptake potential of conjugal plasmids ^60–62^. Further studies are required to confirm if other non-antibiotic pharmaceuticals follow this pattern of enhancing intracellular ROS generation and potentially contributing to increased bacterial gene transfer. Possibly, a ROS measurement in bacteria could be used to screen for non-antibiotic pharmaceuticals that contribute to spreading antibiotic resistance.

We looked for chemical structures and properties of these non-antibiotic pharmaceuticals that might be in common in various antibiotics. Four of the pharmaceuticals, ibuprofen, naproxen, gemfibrozil, and diclofenac, harbour benzene rings and carboxyl functional groups. This is similar to antibiotics such as ampicillin, cefalexin and ciprofloxacin (Fig. S4). A simple chemical comprising a benzene ring and a carboxyl group is salicylic acid. This has been widely demonstrated to behave like an antibiotic on both Gram-positive and Gram-negative bacteria. This includes reducing susceptibility towards antimicrobials ^63, 64^ and inducing intrinsic multiple antibiotic resistance ^65^. In addition, *in vitro* experiments show carboxyl functionalized graphene causes structural damage on plasma membrane and induces intracellular ROS generation at concentration as low as 4 *µ*g/mL ^66^. Carboxyl functionalized graphene also shows toxicity towards *Caenorhabditis elegans* and enhances ROS production *in vivo* ^67^. Therefore, we infer that non-antibiotic pharmaceuticals containing a benzene ring and a carboxyl group may endow them to exhibit antibiotic-like features by enhancing the production of ROS and facilitating the conjugative transfer of ARGs.

This study expands our understanding towards the spread of antibiotic resistance. It is apparent that, in addition to antibiotics, non-antibiotic human-targeted pharmaceuticals will also contribute to the horizontal transfer of ARGs. These findings add to the increasingly complicated nature of the spread of antibiotic resistance. In addition, our findings suggest the antibiotic-like roles of non-antibiotic pharmaceuticals should be considered for pharmaceutical development. Further *in vivo* studies could be conducted to test whether non-antibiotic human-targeted pharmaceuticals facilitate antibiotic resistant bacteria propagation in relevant environments such as in the human gut.

## Materials and Methods

### Bacterial strains and MIC determination

*Escherichia coli* K-12 LE392 with RP4 plasmid (resistant to tetracycline, kanamycin and ampicillin) was the donor. *Pseudomonas putida* KT2440 with high resistance towards chloramphenicol was the recipient ^20, 60^. Culture conditions are described in Text S1. MICs of bacterial strains towards antibiotics and non-antibiotic pharmaceuticals were determined according to previous methods. MICs were calculated based on the comparison between pharmaceutical-dosed groups and the relevant solvent groups, either sterilized MilliQ water or ethanol. Details are described in Text S2 ^20, 68^.

### Conjugative transfer and reverse transfer with the addition of non-antibiotic pharmaceuticals

Both donor and recipient at the concentration of 10^8^ cfu/mL were mixed well at a ratio of 1:1 to establish the PBS-based conjugative mating system (pH=7.2), with a total volume of 1 mL for each mating system. Different levels of non-antibiotic pharmaceuticals were added to the mating system. This included clinical and environmental relevant concentrations, and sub-MIC levels. These were 0.005, 0.05, 0.5, 5, 50 mg/L for ibuprofen, naproxen, gemfibrozil, diclofenac, propanolol, and 0.0001, 0.001, 0.01, 0.1, 1, 5 mg/L for iopromide (due to the solubility). After 8 h-incubation at 25 °C without shaking, 50 *µ*L of the mixture was spread on to LB agar selection plates containing antibiotics to count the number of transconjugants, details are described in Text S3. In addition to the above matings, further sets of conjugative mating systems were established with the addition of 100 *µ*M ROS scavenger, thiourea. The conjugative transfer frequency was calculated from the number of transconjugant colonies divided by the number of recipients. As no nutrient was provided during the mating process, the growths of donor, recipient, and transconjugant were neglected.

To test for the reverse transfer process, transconjugants obtained from transfer experiment were applied as the new donor, while a mutant strain of *E. coli* MG1655 with chloramphenicol resistance was the recipient ^21^. The conjugation experiments were conducted with the different non-antibiotic pharmaceuticals as described above. The number of transconjugants were counted on Difco^TM^ m Endo Agar plates (to distinguish *E. coli* and *P. putida*) with the appropriate antibiotics as described in Text S3.

### Plasmid verification

Transconjugants growing on the selective plates were randomly picked, cultured, and stored with 25% glycerol in −80 °C. The plasmids of transconjugants were extracted using the Invitrogen^TM^ PureLink^®^ Quick Plasmid Miniprep Kit (Life Technologies, USA). The specific *traF* gene of RP4 plasmid was amplified by PCR, and the amplicons were observed using 1% agarose gel electrophoresis. To further verify the identity of the plasmid, PCR was applied for detection of the *tetA* and *bla* genes, which are harboured on the RP4 plasmid. PCR primers and conditions are described in Text S4 and Table S21.

### Transmission electron microscopy

TEM was performed to reveal the influence pharmaceuticals on bacterial cells. Conjugation experiments were performed as described above and TEM samples were collected after 8-h’s mating with either 0.5 mg/L ibuprofen, naproxen, gemfibrozil, diclofenac, propanolol, or 0.1 mg/L iopromide. Sample preparations were performed according to standard procedures as previously described ^69^, and details are illustrated in Text S5. A JEOL JEM-1011 (JEOL, Japan) operated at 80 kV was applied to obtain the images.

### ROS generation and cell membrane permeability detection

ROS generation and cell membrane permeability were detected based on the fluorescence-method as described in Text S6. In brief, 20 *µ*M of DCFDA and 2 mM of propidium iodide (PI) were applied to dye the donor and recipient cells after exposure to the various non-antibiotic pharmaceuticals. The dyed cells were then detected by a CytoFLEX S flow cytometer (Beckman Coulter, USA). The DCFDA- and PI-stained cells were recorded, and calculated as fold changes comparing to the control group (absence of added pharmaceuticals).

### Whole-genome RNA sequence analysis and bioinformatics

In order to analyze the gene expression levels during the conjugative process, the same conjugation experiments were performed as described above and RNA was extracted after 2-h’s mating with either 0.5 mg/L ibuprofen, naproxen, gemfibrozil, diclofenac, propanolol, or 0.1 mg/L iopromide. As bacterial mRNA expressions respond quickly to external stress, 2-h’s mating time was chosen as done previously ^47, 60^. Total RNA (containing the mixture of donor and recipient bacteria) was extracted using RNeasy Mini Kit (QIAGEN^®^, Germany) with an extra bead-beating step for the cell lysis process ^20^. The RNA samples with biological triplicates were then submitted to Macrogen Co. (Seoul, Korea) for strand specific cDNA library construction and Illumina paired-end sequencing (HiSeq 2500, Illumina Inc., San Diego, CA). Raw data were analyzed using the bioinformatic pipeline described previously^69^. Noticeably, the database used for alignment was the combination of reference genome of *E. coli* K-12 (NC_000913), *P. putida* KT2440 (NC_002947), and IncP*α* RP4 plasmid (L27758), which were obtained from National Center for Biotechnology Information (NCBI). Regarding the bioinformatic pipeline, NGS QC Toolkit (v2.3.3), SeqAlto (version 0.5), and Cufflinks (version 2.2.1) were applied to treat the raw sequence reads and to analyze the differential expression for triplicated samples. CummeRbund package in R was used to conduct the statistical analyses. We used the measure of “fragments per kilobase of a gene per million mapped reads” (FPKM) to quantify gene expression. The differences of gene expression between the control (no added pharmaceuticals) and the pharmaceutical-exposed groups were presented as log_2_ fold-changes (LFC).

### Proteomic analysis and bioinformatics

Conjugation experiments were established as described above to compare proteins expressed in the donor and recipient bacteria during the absence and presence of the non-antibiotic pharmaceuticals. Initially, the optimal length of exposure period was examined in the conjugations when exposed to either 0.5 mg/L gemfibrozil or propranolol. Total proteins from the mixture of donor and recipient bacteria were extracted after 2, 4, 6, and 8 h mating as described previously ^20^. For peptide preparations, the extracted proteins were treated by reduction, alkylation, trypsin digestion, and ziptip clean-up procedures as described previously ^70^. The peptide preparations were then loaded to mass spectrometer. Qualitative protein libraries were constructed by information dependent analysis; while quantitative protein determination was based on SWATH-MS ^70^ using biological triplicate samples. Database and software analyses and settings were performed as described in Text S7. A stringency cut-off of false discovery rate (*q* value) less than 0.01 was used to identify the proteins with significant different expression levels. Based on the number of proteins showing significant variations, 8 h was determined as the best exposure time for the proteomic analysis. Thus, another set of conjugation experiment was established as described above using the 8 h mating period in the presence of either ibuprofen, naproxen, gemfibrozil, diclofenac, or propranolol, each at 0.5 mg/L, or with iopromide at 0.1 mg/L. Following that, for each of the conjugation experiments, the proteins were extracted, peptide preparations prepared and proteomic analyses were performed as described above.

### Correlation tests

Correlation tests were conducted to identify whether the phenotypic data (including conjugative transfer frequency, ROS generation and cell membrane permeability) were concentration-dependent. Pearson correlation formula (Eq. 1) was applied to calculate the correlation coefficient value *r*, followed by consulting the correlation coefficient table. The correlation was significant if *P* values were less than 0.05.

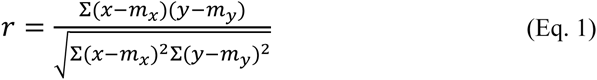

### Statistical analysis

Data were expressed as mean ± standard deviation (SD). SPSS for Mac version 25.0 was applied for data analysis. Independent-sample *t* tests were performed. *P* values less than 0.05 were considered to be statistically significant. All the experiments were conducted in triplicate.

## Data availability

All data was deposited in publicly accessible databases. RNA sequence data are accessible through Gene Expression Omnibus of NCBI (GSE130562).The mass spectrometry proteomics data have been deposited to the ProteomeXchange Consortium via the PRIDE ^71^ partner repository with the dataset identifier of PXD012642.

## Acknowledgements

We acknowledge the Australian Research Council for funding support through Future Fellowship (FT170100196). Jianhua Guo would like to thank the support by UQ Foundation Research Excellence Awards. Yue Wang would like to thank the support from China Scholarship Council. We thank Prof. Mark Walker (The University of Queensland) for providing *E. coli* with RP4 plasmid. We would like to thank Dr. Michael Nefedov (The University of Queensland) for providing technical support on flow cytometry. We would also like to thank Dr. Amanda Nouwens (The University of Queensland) for conducting SWATH-MS tests. The MIC measurement in this work was performed at the Queensland node of the Australian National Fabrication Facility.

## Author Contributions

J.G. and Y.W. conceived and designed this study; Y.W. performed the sampling, transfer experiment, flow cytometer, and DNA, RNA and protein extractions. J.L. conducted the transmission electron microscopy testing. S.Z. performed the RNA extraction. L.M. and J.L. analyzed transcriptomic data. J.G. Y.W. and P.B provided critical biological interpretations of the data. Y.W. and J.G. wrote the manuscript. J.G., P.B. and Z.Y. supervised this work and edited on the manuscript.

