## Supplemental Information for "Non-antibiotic pharmaceuticals can enhance the spread of antibiotic resistance via conjugation"

**This file includes:**

Supplementary Methods 1 to 7

Supplementary Figures 1 to 4

Supplementary Tables 1 to 21

#### Supplementary Methods

##### Text S1. Culture conditions for donor and recipient bacteria

Both donor and recipient were cultured separately in Luria-Bertani (LB) broth (pH 7.0) at 30 $^{o}$C for 16 h with the supplementary of appropriate antibiotics. For donor, 17.0 mg/L tetracycline, 33.0 mg/L kanamycin, and 100.0 mg/L ampicillin were added, while 17.0 mg/L chloramphenicol was dosed in the LB broth for recipient. After culturing, both donor and recipient were washed with phosphate-buffered saline (PBS, pH=7.2) twice to eliminate the possible influence induced by culture media. Afterwards, the donor and recipient were re-suspended separately in different volumes of PBS to obtain initial concentration of $10^8$ cfu/mL based on OD600 values. Then, the donor and recipient were mixed with the ratio of 1:1. The mixtures were applied immediately for the conjugation experiment.

##### Text S2. Determination of MICs

The bacteria with initial concentration of $10^5$ cfu/mL were used for MIC detection. In each well of the 96-well plates, 5 $\mu$L of the bacteria, 15 $\mu$L of antibiotics or non-antibiotic pharmaceuticals with different concentrations, and 130 $\mu$L of LB was added. Blank controls were ethanol or sterilized MilliQ water. The 96-well plates were then incubated at 30 $^{o}$C for 18 h, followed by OD600 measured on the plate reader (Tecan Infinite M200, Switzerland). MICs were determined as the concentration of antibiotics or non-antibiotic pharmaceuticals that could inhibit at least 90% of bacterial growth. MICs were tested in triplicate.

##### Text S3. Selection plates for transconjugant and recipient

The selection plates for transconjugant contained all of the four kinds of antibiotics (17.0 mg/L tetracycline, 33.0 mg/L kanamycin, and 100.0 mg/L ampicillin), while those for recipient only contained 17.0 mg/L chloramphenicol. After plating on transconjugant and recipient selective plates, the plates were incubated at 30 $^{o}$C for 48 h, followed by counting the colonies growing on them. The results of selection plates are shown as cfu/mL, and transfer frequency was calculated as the number of transconjugants divided by the number of recipients. All the selection plates were performed at least in triplicate. In addition, both donor and recipient were plated onto the transconjugant selective plates, to rule out any spontaneous mutation.

**Text S4. PCR conditions**

PCR systems were set up as 25 $\mu$L, with 12.5 $\mu$L Platinum™ Green Hot Start PCR Master Mix (2X) (Invitrogen$^{TM}$), 1 $\mu$L 20 $\mu$M primer, 1 $\mu$L plasmid, and 10.5 $\mu$L dd$H_2O$. Primers are listed in Supplementary table 21. PCR conditions for gene *traF* were: denaturation at 94 $^{o}$C for 4 min on initial cycle, 30 s for another 35 cycles, annealing at 55 $^{o}$C for 30 s, extension at 72 $^{o}$C for 1 min, followed by 7 min. The process was conducted with 30 cycles [1]. PCR conditions for gene *tetA* and *bla* were the same as those for gene *traF*, except with the annealing temperature of 54 $^{o}$C, and 50 $^{o}$C respectively.

**Text S5. Sample preparations for transmission electron microscopy**

Cells were collected by centrifuging at 4000 g for 5 min, followed by fixing in PBS with 2.5% glutaraldehyde and stored at 4 °C overnight. Afterwards, cell pellets were re-suspended in PBS and microwaved twice at 80 W for 40 s. Following centrifuge, the cell pellets were heated to 37 $^{o}$C and soaked with 2% agarose. The cell pellets were then solidified, cut into small pieces, added with 1% osmium tetroxide, and microwaved twice at 80 W for 2 min. Osmium tetroxide was then discarded. Samples were further processed by dehydration, infiltration and mounted with Epon and polymerized at 60 °C for 2 days [2]. Ultrathin sections (50–100 nm) of samples were then collected on TEM grid and observed using a 80 kV JEOL JEM-1011 (JEOL, Japan).

**Text S6. ROS generation and cell membrane permeability detection**

Bacteria strains were washed twice with PBS and resuspended in PBS to $10^6$ cfu/mL. For ROS detection, bacteria strains were incubated in dark at 37 $^{o}$C for 30 min with 2', 7'-dichlorofluorescein diacetate (DCFDA, at a final concentration of 20 uM, abcam®). Then, 100 µL of the bacteria stained with DCFDA were treated with different concentrations of non-antibiotic pharmaceuticals. 1.5% $H_2O_2$ was set as positive control, and ethanol was set as negative control. After complete mixing by vortex, the mixtures were incubated in dark at 25 $^{o}$C for 2 h before measurement at 488 nm. For cell membrane permeability detection, 100 µL of bacteria strain was exposed to different concentrations of non-antibiotic pharmaceuticals, and incubated at 25 $^{o}$C for 2 h. The same volume of ethanol was the negative control, while bacteria strain treated with 100 $^{o}$C water was the positive control. The strains were then stained with 1 µL of propidium iodide (PI, 2 mM, Life Technologies) and incubated in the dark for 15 min before measurement at 561 nm. All data was analysed with CytExpert. Data were presented in a dot plot, in which upper right quadrant indicated DCFDA or PI positive cells (with increased ROS or cell membrane permeability), and upper left quadrant was normal cells. All the detections were conducted in triplicate. Relative fold increases in ROS production or cell membrane permeability were calculated as pharmaceutical-treated samples divided by negative control samples according to previous studies [3,4].

**Text S7. Proteomics analysis**

Qualitative protein libraries were constructed by information dependent analysis (IDA), and quantitative protein determination was based on SWATH-MS. IDA data were combined and searched using ProteinPilot software, with the combined databases of *E. coli* SP only (received from Uniprot on 9$^{th}$ of July 2018) and *P. putida* KT2440 (received from NCBI on 9$^{th}$ of July 2018). Search setting for enzyme digestion was set to trypsin and alkylation was set to iodoacetamide. Afterwards, the constructed IDA library and SWATH-MS data were loaded into PeakView v2.1 for further processing, with the peptide confidence threshold of 99%, number of peptides per protein of 5, and number of transitions per peptide of 3. A minimum of 2 peptides and 3 transitions was used for quantitative analysis.

### Supplementary Figures

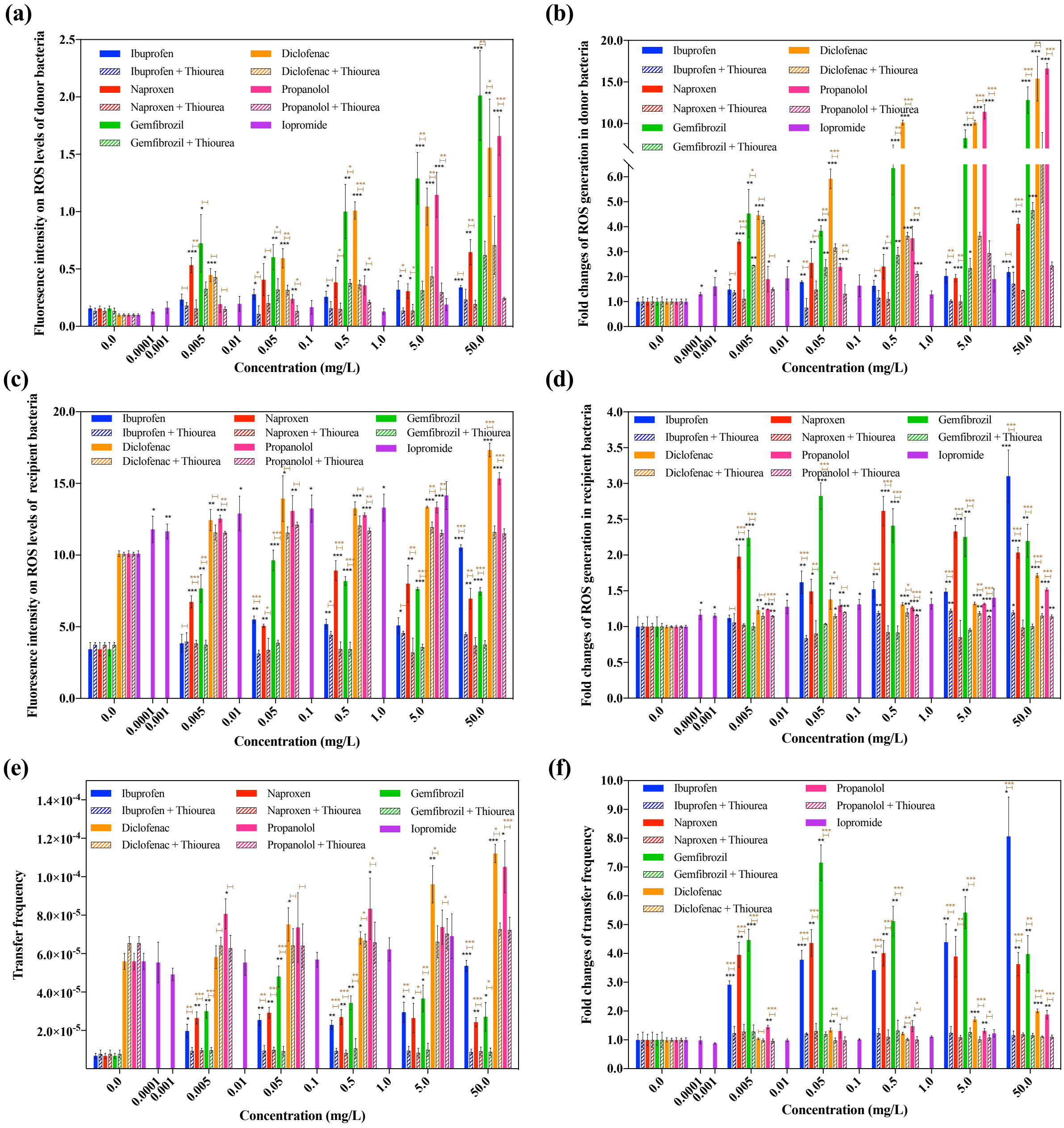


Fig. S1. Effects of non-antibiotic pharmaceuticals and thiourea on ROS in the donor (*E. coli* K-12 LE392) and recipient (*P. putida* KT2440) bacteria. (a) Fluorescence intensity on ROS levels of donor bacteria. (b) Fold changes of ROS generation in donor bacteria. (c) Fluorescence intensity on ROS levels of recipient bacteria. (d) Fold changes of ROS generation in recipient bacteria. (e) Conjugative transfer frequency with the addition of ROS scavenger thiourea. (f) Fold changes of conjugative transfer frequency with the addition of ROS scavenger thiourea. Significant differences between non-antibiotic-dosed samples and the control were analyzed by independent-sample *t* test, *$P$<0.05, **$P$<0.01, and ***$P$<0.001.

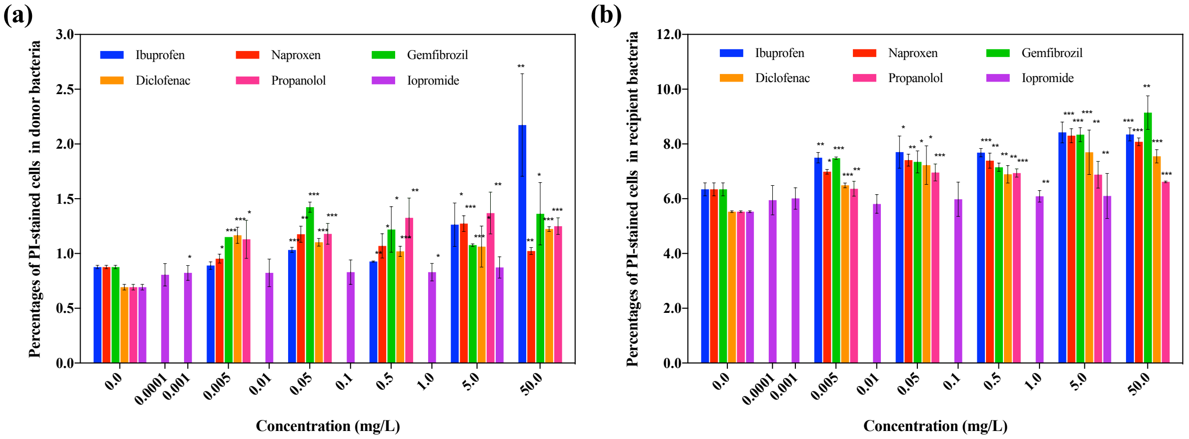


Fig. S2. Effects of non-antibiotic pharmaceuticals on cell membranes in the donor (E. coli K-12 LE392) and recipient (P. putida KT2440) bacteria. (a) Percentages of PI-stained cells in donor bacteria. (b) Percentages of PI-stained cells in recipient bacteria. Significant differences between non-antibiotic-dosed samples and the control were analyzed by independent-sample *t* test, **P*<0.05, ***P*<0.01, and ****P*<0.001.

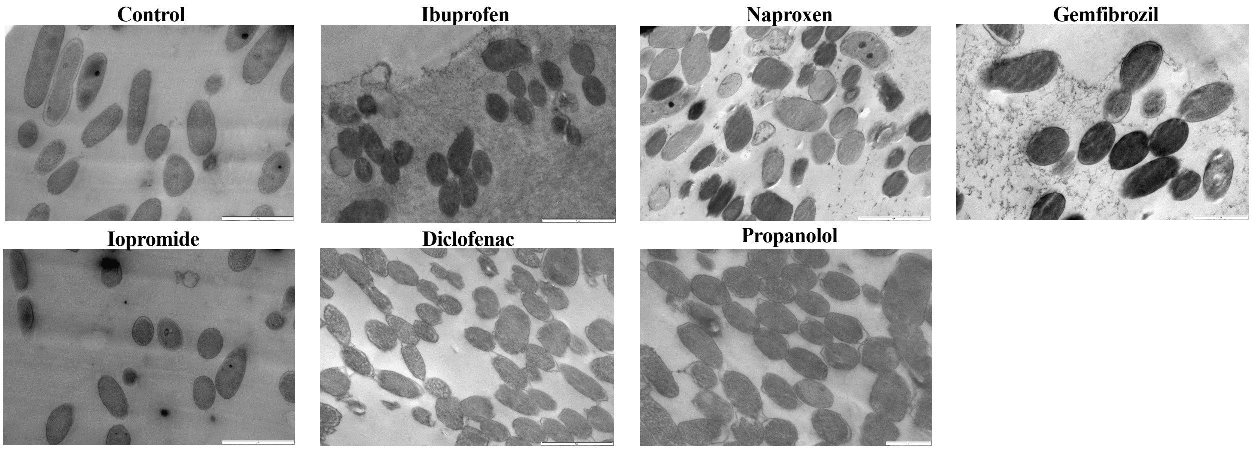


Fig. S3. TEM images of donor and recipient bacteria under the exposure of non-antibiotic pharmaceuticals.

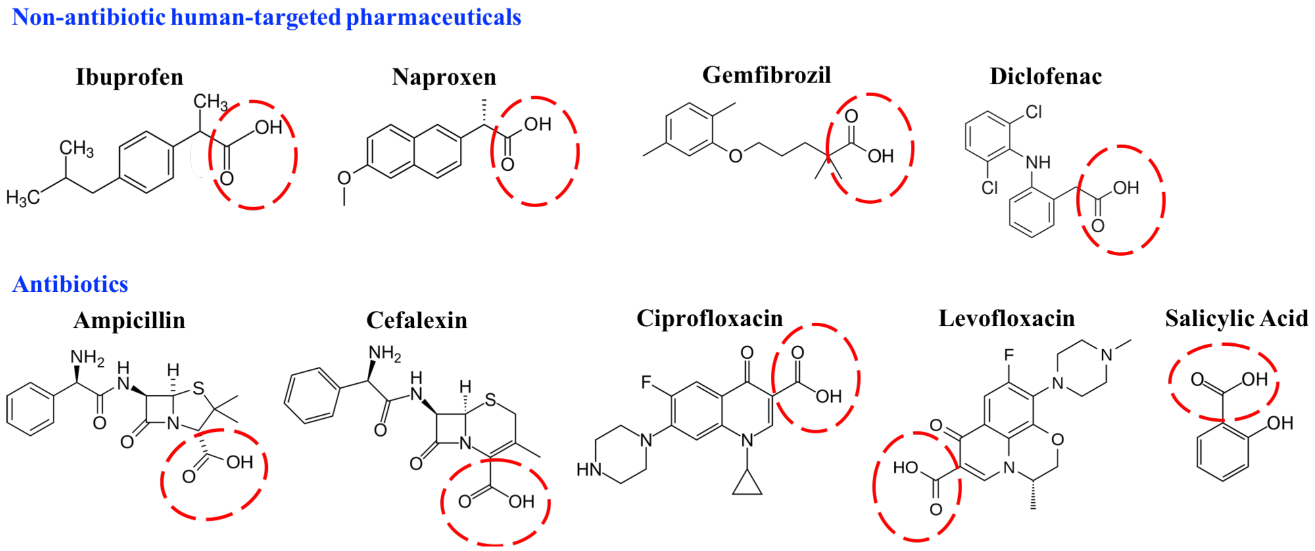


Fig. S4. Chemical structures of non-antibiotic human-targeted pharmaceuticals that can promote conjugation, and chemical structures of commonly prescribed antibiotics. All of these drugs harbour benzene ring and carboxyl functional group.

**Supplementary Tables**

Table S1. Minimum inhibitory concentrations (MICs) of donor and recipient bacterial strains towards non-antibiotic pharmaceuticals

| Strains | MICs (mg/L) | | | | | |
|---|---|---|---|---|---|---|
| | Ibuprofen | Naproxen | Gemfibrozil | Iopromide | Diclofenac | Propanolol |
| Donor (*E. coli* LE392) | 500 | 500 | >500 | >10 | >500 | 500 |
| Recipient (*P. putida* KT2440) | 500 | 500 | >500 | >50 | >1000 | 500 |

Table S2. Minimum inhibitory concentrations (MICs) of donor, recipient, and different transconjugants towards antibiotics*

| Antibiotics | MICs (mg/L) | | | | | | | | | |
|---|---|---|---|---|---|---|---|---|---|---|
| | Donor | Recipient | TC 1 | TC 2 | TC 3 | TC 4 | TC 5 | TC 6 | TC 7 | TC 8 |
| Tetracycline | 12.5 | 1.25 | 12.5 | 12.5 | 12.5 | 12.5 | 12.5 | 12.5 | 12.5 | 12.5 |
| Kanamycin | 20 | 5 | 20 | 20 | 20 | 20 | 20 | 20 | 20 | 20 |
| Ampicillin | >120 | 6 | >120 | >120 | >120 | >120 | >120 | >120 | >120 | >120 |
| Chloram-phenicol | 0.13 | 0.39 | 0.39 | 0.39 | 0.39 | 0.39 | 0.39 | 0.39 | 0.39 | 0.39 |

* TC 1-8: transconjugants in mating system treated with Milli-Q water, ethanol, ibuprofen, naproxen, gemfibrozil, iopromide, diclofenac, propanolol, respectively

Table S3. Genes relevant to ROS production in donor bacteria *E. coli* K-12 LE392 after exposure of non-antibiotic pharmaceuticals

| **Gene** | **COG Annotation** | **Fold Change of FPKM** * | | | | | |
|---|---|---|---|---|---|---|---|
| | | **Ibuprofen** | **Naproxen** | **Gemfibrozil** | **Diclofenac** | **Propanolol** | **Iopromide** |
| *ahpC* | Alkyl hydroperoxide reductase subunit AhpC (peroxiredoxin) | 1.44 | 1.33 | 1.04 | 1.39 | 1.37 | 1.32 |
| *ahpF* | Alkyl hydroperoxide reductase subunit AhpF | 1.07 | 0.94 | 0.85 | 1.06 | 1.04 | 1.13 |
| *alkB* | Alkylated DNA repair dioxygenase AlkB | 1.19 | 4.79 | 1.69 | 1.60 | 2.14 | 1.56 |
| *oxyR* | DNA-binding transcriptional regulator, LysR family | 1.51 | 1.31 | 1.04 | 1.27 | 1.15 | 1.06 |
| *rutC* | Enamine deaminase RidA, house cleaning of reactive enamine intermediates, YjgF/YER057c/UK114 family | 1.34 | 1.48 | 10.85 | 1.64 | 1.35 | 1.11 |
| *rutE* | Homoserine acetyltransferase | 2.23 | 10.56 | 1.29 | 1.65 | 1.26 | 1.31 |
| *sodB* | Superoxide dismutase | 1.55 | 1.40 | 1.03 | 1.03 | 1.20 | 1.19 |
| *sodC* | Superoxide dismutase | 1.46 | 2.27 | 1.01 | 1.01 | 1.22 | 1.01 |

| Gene | COG Annotation | Fold Change of FPKM * | | | | | |
|---|---|---|---|---|---|---|---|
| | | **Ibuprofen** | **Naproxen** | **Gemfibrozil** | **Diclofenac** | **Propanolol** | **Iopromide** |
| *soxR* | DNA-binding transcriptional regulator, MerR family | 1.49 | 1.08 | 1.09 | 1.10 | 1.57 | 1.16 |
| *soxS* | AraC-type DNA-binding domain and AraC-containing proteins | 1.25 | 1.16 | 1.04 | 1.27 | 0.95 | 1.03 |
| *trxC* | Negative regulator of GroEL, contains thioredoxin-like and TPR-like domains | 2.77 | 2.06 | 1.25 | 2.00 | 2.81 | 0.94 |

*: Comparing with the control group without pharmaceutical dosage

Table S4. Proteins relevant to ROS production in donor bacteria *E. coli* K-12 LE392 after exposure of non-antibiotic pharmaceuticals

| Protein | Description | Fold Change of Protein Abundance * | | | | | |
|---|---|---|---|---|---|---|---|
| | | **Ibuprofen** | **Naproxen** | **Gemfibrozil** | **Diclofenac** | **Propanolol** | **Iopromide** |
| AhpF | Alkyl hydroperoxide reductase subunit F | 0.93 | 1.02 | 1.06 | 1.08 | 1.12 | 1.05 |
| SodC | Superoxide dismutase | 2.17 | 2.25 | 1.46 | 2.30 | 4.72 | 2.27 |

*: Comparing with the control group without pharmaceutical dosage

Table S5. Genes relevant to ROS production in recipient bacteria *P. putida* KT2440 after exposure of non-antibiotic pharmaceuticals

| **Gene** | **COG Annotation** | **Fold Change of FPKM** * | | | | | |
|---|---|---|---|---|---|---|---|
| | | **Ibuprofen** | **Naproxen** | **Gemfibrozil** | **Diclofenac** | **Propanolol** | **Iopromide** |
| *oxyR* | Oxidative and nitrosative stress transcriptional dual regulator | 1.54 | 1.27 | 1.17 | 1.14 | 1.11 | 1.04 |
| *sodA* | Superoxide dismutase | 1.40 | 2.07 | 3.32 | 1.37 | 2.66 | 1.34 |
| *sodB* | Superoxide dismutase | 1.17 | 1.21 | 1.02 | 1.11 | 1.05 | 0.95 |

*: Comparing with the control group without pharmaceutical dosage

Table S6. Proteins relevant to ROS production in recipient bacteria *P. putida* KT2440 after exposure of non-antibiotic pharmaceuticals

| **Protein** | **Description** | **Fold Change of Protein Abundance** * | | | | | |
|---|---|---|---|---|---|---|---|
| | | **Ibuprofen** | **Naproxen** | **Gemfibrozil** | **Diclofenac** | **Propanolol** | **Iopromide** |
| AhpC | Alkyl hydroperoxide reductase subunit C | 2.73 | 1.73 | 1.87 | 1.31 | 1.08 | 1.95 |
| SodF | Superoxide dismutase | 1.07 | 0.91 | 0.99 | 0.95 | 0.62 | 0.76 |
| Tpx | Thioredoxin peroxidase | 1.40 | 1.08 | 1.12 | 0.90 | 0.95 | 1.21 |

*: Comparing with the control group without pharmaceutical dosage

Table S7. Genes relevant to cell membrane in donor bacteria *E. coli* K-12 LE392 after exposure of non-antibiotic pharmaceuticals

| Gene | COG Annotation | Fold Change of FPKM * | | | | | |
|---|---|---|---|---|---|---|---|
| | | Ibuprofen | Naproxen | Gemfibrozil | Diclofenac | Propanolol | Iopromide |
| *csgG* | Curli production assembly/transport outer membrane lipoprotein | 3.73 | 4.72 | 1.54 | 3.05 | 1.77 | 1.13 |
| *cusA* | Efflux system membrane component | 1.25 | 1.26 | 2.07 | 0.75 | 1.01 | 0.67 |
| *ompC* | Outer membrane porin protein C | 1.19 | 1.08 | 1.24 | 1.07 | 1.12 | 0.94 |
| *ompF* | Outer membrane porin 1a | 2.51 | 2.38 | 1.02 | 1.82 | 2.00 | 1.04 |
| *ompN* | Outer membrane pore protein non-specific | 1.11 | 1.32 | 1.39 | 2.35 | 1.69 | 1.04 |
| *ompR* | Response regulator in two-component regulatory system | 1.03 | 1.23 | 1.01 | 1.37 | 1.35 | 1.51 |
| *ompT* | Outer membrane protease | 1.45 | 1.21 | 0.35 | 1.21 | 1.26 | 0.61 |
| *ompW* | Outer membrane protein W | 0.70 | 1.12 | 1.74 | 1.92 | 1.18 | 0.86 |

| Gene | COG Annotation | Fold Change of FPKM [*] | | | | | |
|---|---|---|---|---|---|---|---|
| | | Ibuprofen | Naproxen | Gemfibrozil | Diclofenac | Propanolol | Iopromide |
| *pgaA* | Biofilm adhesin polysaccharide PGA secretin OM porin export protein | 1.80 | 2.55 | 2.93 | 2.75 | 2.25 | 1.27 |
| *ybhG* | Putative membrane fusion protein (MFP) component of efflux pump membrane anchor | 1.16 | 1.14 | 4.72 | 1.80 | 2.33 | 2.97 |
| *ydcU* | Putative ABC transporter permease | 7.26 | 4.20 | 2.36 | 1.61 | 7.94 | 1.09 |
| *yfaZ* | Outer membrane protein putative porin | 1.53 | 1.78 | 1.31 | 1.31 | 1.51 | 1.11 |

[*]: Comparing with the control group without pharmaceutical dosage

Table S8. Proteins relevant to cell membrane in donor bacteria *E. coli* K-12 LE392 after exposure of non-antibiotic pharmaceuticals

| Protein | Description | Fold Change of Protein Abundance * | | | | | |
|---|---|---|---|---|---|---|---|
| | | **Ibuprofen** | **Naproxen** | **Gemfibrozil** | **Diclofenac** | **Propanolol** | **Iopromide** |
| BamB | Outer membrane protein assembly factor | 0.77 | 0.93 | 0.96 | 1.06 | 1.21 | 1.19 |
| OmpC | Outer membrane protein C | 1.28 | 1.54 | 1.59 | 1.51 | 2.16 | 2.06 |
| OmpF | Outer membrane protein F | 1.61 | 1.52 | 1.20 | 1.57 | 1.62 | 2.30 |
| Slp | Outer membrane protein | 0.72 | 0.84 | 0.83 | 1.06 | 0.99 | 0.81 |

*: Comparing with the control group without pharmaceutical dosage

Table S9. Genes relevant to cell membrane in recipient bacteria *P. putida* KT2440 after exposure of non-antibiotic pharmaceuticals

| **Gene** | **COG Annotation** | **Fold Change of FPKM** * | | | | | |
|---|---|---|---|---|---|---|---|
| | | **Ibuprofen** | **Naproxen** | **Gemfibrozil** | **Diclofenac** | **Propanolol** | **Iopromide** |
| *czcB-I* | Function of homologous gene experimentally demonstrated in another organism Product type m: membrane componentTransport and binding proteins | 1.49 | 3.86 | 1.93 | 1.33 | 1.06 | 1.83 |
| *czcC* | RND transporter outer membrane protein | 1.39 | 1.46 | 1.40 | 0.83 | 2.16 | 0.64 |
| *ompQ* | outer membrane pyoverdine efflux protein | 1.39 | 1.73 | 1.40 | 1.46 | 1.75 | 0.74 |
| *ompR* | two-component system DNA-binding response regulator | 1.82 | 1.52 | 1.25 | 0.96 | 0.92 | 0.76 |
| *opdT-II* | Tyrosine-specific outer membrane porin D | 1.68 | 1.55 | 1.73 | 1.11 | 1.31 | 1.11 |
| *oprG* | Outer membrane protein OprG | 0.99 | 0.99 | 1.27 | 1.32 | 1.29 | 1.31 |
| *oprH* | Outer membrane protein H1 | 1.24 | 1.18 | 1.09 | 1.16 | 1.21 | 1.24 |

| Gene | COG Annotation | Fold Change of FPKM * | | | | | |
|---|---|---|---|---|---|---|---|
| | | Ibuprofen | Naproxen | Gemfibrozil | Diclofenac | Propanolol | Iopromide |
| *oprI* | Major outer membrane lipoprotein | 1.16 | 0.99 | 1.48 | 1.13 | 1.24 | 1.34 |
| *oprJ* | Outer membrane protein OprJ | 1.21 | 1.97 | 1.36 | 1.15 | 1.39 | 1.12 |
| *PP_0143* | Membrane protein | 0.85 | 1.18 | 1.27 | 1.34 | 1.35 | 0.86 |
| *PP_0426* | Membrane protein | 1.06 | 0.88 | 1.18 | 1.27 | 1.06 | 1.06 |
| *PP_0717* | Membrane protein | 1.46 | 1.84 | 3.34 | 1.56 | 1.02 | 0.72 |
| *PP_0828* | Membrane protein | 2.17 | 1.78 | 1.45 | 0.88 | 2.95 | 1.89 |
| *PP_0984* | Membrane protein | 1.91 | 1.83 | 1.19 | 1.39 | 1.84 | 2.00 |
| *PP_1150* | Membrane protein | 1.80 | 1.62 | 1.34 | 1.37 | 1.16 | 1.01 |
| *PP_1159* | Membrane protein | 2.16 | 0.58 | 1.39 | 1.59 | 0.74 | 1.38 |
| *PP_1359* | Membrane protein | 1.15 | 1.13 | 2.85 | 1.88 | 2.17 | 1.16 |
| *PP_1728* | Membrane protein | 1.42 | 1.35 | 0.83 | 1.44 | 1.01 | 0.59 |
| *PP_1936* | Membrane protein | 1.51 | 1.66 | 1.57 | 1.35 | 1.97 | 1.22 |
| *PP_2014* | Membrane protein | 1.47 | 1.12 | 1.83 | 1.22 | 1.01 | 1.09 |
| *PP_2104* | Membrane protein | 0.93 | 1.16 | 1.51 | 1.08 | 1.07 | 1.16 |

| Gene | COG Annotation | Fold Change of FPKM * | | | | | |
|---|---|---|---|---|---|---|---|
| | | Ibuprofen | Naproxen | Gemfibrozil | Diclofenac | Propanolol | Iopromide |
| *PP_2384* | Membrane protein | 2.19 | 2.31 | -- | 3.56 | -- | 2.22 |
| *PP_2401* | Membrane protein | 1.27 | 1.06 | -- | 3.16 | 2.08 | -- |
| *PP_2429* | Membrane protein | 0.95 | 1.45 | 1.88 | 1.78 | 2.23 | 1.55 |
| *PP_2721* | Membrane protein | 1.44 | 12.64 | 2.77 | 2.57 | 2.79 | 3.14 |
| *PP_3105* | Membrane protein | 1.40 | 2.33 | 2.38 | 1.17 | 1.39 | 1.68 |
| *PP_3169* | Membrane protein | 1.09 | 1.69 | 1.51 | 1.11 | 0.95 | 2.06 |
| *PP_3329* | Membrane protein | 1.11 | 0.97 | 1.30 | 1.28 | 1.66 | 1.53 |
| *PP_3389* | Membrane protein | 1.91 | 2.17 | 4.96 | 1.68 | 1.30 | 3.23 |
| *PP_3609* | Membrane protein | 1.46 | 1.26 | 2.69 | 1.18 | 1.62 | 0.86 |
| *PP_3661* | Membrane protein | 1.27 | 1.30 | 2.45 | 1.47 | 1.58 | 1.80 |
| *PP_4118* | Membrane protein | 1.24 | 1.34 | 2.25 | 1.72 | 1.16 | 2.33 |
| *PP_4598* | Membrane protein | 1.01 | 1.19 | 1.60 | 1.95 | 1.74 | 2.10 |
| *PP_4771* | Membrane protein | 2.75 | 3.27 | 1.91 | 1.09 | 0.69 | 0.57 |
| *PP_4815* | Membrane protein | 1.04 | 1.32 | 1.37 | 1.23 | 1.15 | 1.46 |

| Gene | COG Annotation | Fold Change of FPKM * | | | | | |
|---|---|---|---|---|---|---|---|
| | | Ibuprofen | Naproxen | Gemfibrozil | Diclofenac | Propanolol | Iopromide |
| *PP_4954* | Membrane protein | 0.39 | 0.03 | 0.52 | 0.22 | -0.11 | 0.33 |
| *PP_5091* | Membrane protein | 0.07 | 0.58 | 1.78 | 1.11 | 1.22 | 1.62 |
| *PP_5133* | Membrane protein | 1.39 | -0.27 | 0.88 | 0.33 | -0.28 | 0.06 |
| *PP_5460* | Membrane protein | 0.76 | 0.43 | 1.08 | 0.36 | -0.25 | 0.47 |

*: Comparing with the control group without pharmaceutical dosage

--: not detected

Table S10. Proteins relevant to cell membrane in recipient bacteria *P. putida* KT2440 after exposure of non-antibiotic pharmaceuticals

| Protein | Description | Fold Change of Protein Abundance * | | | | | |
|---|---|---|---|---|---|---|---|
| | | Ibuprofen | Naproxen | Gemfibrozil | Diclofenac | Propanolol | Iopromide |
| BamA | Outer membrane protein assembly factor | 2.23 | 1.15 | 1.78 | 1.59 | 1.49 | 1.95 |
| OmpA | OmpA family protein | 1.55 | 1.28 | 1.78 | 2.04 | 1.11 | 1.39 |
| OprD | Basic amino acid specific porin | 1.59 | 1.09 | 1.23 | 1.01 | 0.79 | 1.05 |
| OprE | Outer-membrane porin E | 1.35 | 0.91 | 1.13 | 0.79 | 0.90 | 1.06 |
| OprG | Outer membrane protein OprG | 2.41 | 1.75 | 2.00 | 1.21 | 0.84 | 1.30 |
| OprH | Outer membrane protein H1 | 1.39 | 0.97 | 1.24 | 1.17 | 1.09 | 1.08 |
| OprI | Major outer membrane lipoprotein | 1.24 | 1.09 | 1.44 | 1.01 | 0.68 | 0.72 |
| OprL | Peptidoglycan-associated lipoprotein | 1.31 | 0.95 | 1.11 | 0.99 | 0.97 | 1.49 |
| OprQ | Outer-membrane porin D | 1.69 | 1.21 | 1.34 | 1.37 | 1.27 | 1.37 |
| TtgC | Probable efflux pump outer membrane protein TtgC | 1.93 | 1.05 | 1.77 | 1.21 | 1.38 | 1.58 |

*: Comparing with the control group without pharmaceutical dosage

Table S11. Genes relevant to conjugative transfer, pilus generation, plasmid replication in IncP-α RP4 plasmid after exposure of non-antibiotic pharmaceuticals

| **Gene** | **COG Annotation** | **Fold Change of FPKM** * | | | | | |
|---|---|---|---|---|---|---|---|
| | | **Ibuprofen** | **Naproxen** | **Gemfibrozil** | **Diclofenac** | **Propanolol** | **Iopromide** |
| *korB* | Global regulator | 0.81 | 0.94 | 0.60 | 0.86 | 0.74 | 0.94 |
| *traG* | Conjugative transfer transcriptional regulator | 2.16 | 1.52 | 0.50 | 1.01 | 0.59 | 0.80 |
| *trbD* | Conjugative transfer transcriptional regulator | 1.72 | 1.88 | 2.11 | 1.09 | 0.65 | 0.60 |
| *trbA* | Mating-pair apparatus | 1.99 | 2.27 | 1.08 | 1.25 | 0.92 | 0.87 |
| *trbK* | Mating-pair apparatus | 3.71 | 3.68 | 2.36 | 0.76 | 3.39 | 0.32 |
| *trfA2* | Mating-pair apparatus | 235.57 | 132.51 | 272.48 | 56.49 | 148.06 | 0.07 |
| *traC1* | Replication regulator | 2.01 | 2.04 | 1.38 | 1.36 | 1.40 | 1.27 |
| *traB* | Pilin regulator | 1.95 | 1.42 | 1.10 | 1.16 | 0.86 | 0.99 |
| *traE* | Pilin regulator | 3.25 | 3.03 | 1.60 | 1.22 | 1.99 | 0.65 |
| *traF* | Pilin regulator | 1.12 | 15.45 | 2.39 | 1.39 | 1.60 | 1.15 |
| *traP* | Pilin regulator | 1.28 | 1.17 | 1.10 | 1.28 | 0.88 | 0.88 |

*: Comparing with the control group without pharmaceutical dosage

Table S12. Genes relevant to fimbriae in donor bacteria *E. coli* K-12 LE392 after exposure of non-antibiotic pharmaceuticals

| Gene | COG Annotation | Fold Change of FPKM * | | | | | |
|---|---|---|---|---|---|---|---|
| | | Ibuprofen | Naproxen | Gemfibrozil | Diclofenac | Propanolol | Iopromide |
| *ecpA* | ECP pilin | 0.70 | 1.78 | 4.72 | 1.69 | 3.43 | 1.83 |
| *fimH* | Minor component of type 1 fimbriae | 1.59 | 1.38 | 2.77 | 1.01 | 1.78 | 2.07 |
| *fliI* | Flagellum-specific ATP synthase | 0.72 | 8.17 | 2.30 | 3.18 | 1.36 | 6.50 |
| *hofC* | Assembly protein in type IV pilin biogenesis transmembrane protein | 0.59 | 1.12 | 3.56 | 1.19 | 1.56 | 5.54 |
| *yadN* | Putative fimbrial-like adhesin protein | 3.34 | 3.71 | 3.20 | 3.36 | 2.87 | 1.55 |
| *ybgO* | Putative fimbrial protein | 0.95 | 2.36 | 1.83 | 1.69 | 1.55 | 0.85 |
| *ybgP* | Putative periplasmic pilin chaperone | 3.14 | 1.54 | 7.46 | 3.29 | 5.35 | 1.06 |
| *ycbV* | Putative fimbrial-like adhesin protein | 3.63 | 4.44 | 2.36 | 0.74 | 6.28 | 0.73 |

| Gene | COG Annotation | Fold Change of FPKM * | | | | | |
|---|---|---|---|---|---|---|---|
| | | **Ibuprofen** | **Naproxen** | **Gemfibrozil** | **Diclofenac** | **Propanolol** | **Iopromide** |
| *yfcQ* | Putative fimbrial-like adhesin protein | 6.36 | 4.72 | 6.50 | 11.63 | 1.21 | 1.21 |
| *yfcS* | Putative periplasmic pilin chaperone | 1.71 | 1.47 | 2.01 | 2.55 | 0.74 | 0.82 |
| *yqiI* | Fimbrial protein | 1.06 | 6.73 | 5.94 | 2.14 | 1.08 | 0.55 |
| *yraH* | Putative fimbrial-like adhesin protein | 1.75 | 1.23 | 4.56 | 4.35 | 1.29 | 1.73 |
| *yraI* | Putative periplasmic pilin chaperone | 1.84 | 1.93 | 17.75 | 2.25 | 0.90 | 0.82 |
| *yraK* | Putative fimbrial-like adhesin protein | 1.06 | 0.59 | 1.22 | 1.44 | 1.65 | 1.66 |

*: Comparing with the control group without pharmaceutical dosage

174 Table S13. Genes relevant to fimbriae in recipient bacteria *P. putida* KT2440 after exposure of non-antibiotic pharmaceuticals

| **Gene** | **COG Annotation** | **Fold Change of FPKM** * | | | | | |
|---|---|---|---|---|---|---|---|
| | | **Ibuprofen** | **Naproxen** | **Gemfibrozil** | **Diclofenac** | **Propanolol** | **Iopromide** |
| *flgA* | Flagella basal body P-ring formation protein | 1.31 | 1.20 | 2.07 | 1.68 | 1.23 | 1.21 |
| *pilE* | Type IV pili biogenesis protein | 1.15 | 1.11 | 2.17 | 1.09 | 2.22 | 2.08 |
| *pilH* | Twitching motility protein | 1.41 | 1.23 | 1.61 | 1.20 | 1.32 | 0.99 |
| *pilI* | Twitching motility protein | 1.30 | 1.05 | 1.07 | 1.89 | 1.66 | 1.56 |
| *pilJ* | Twitching motility protein | 1.45 | 1.15 | 1.66 | 1.16 | 1.40 | 1.24 |
| *pilQ* | Type IV pili biogenesis protein | 1.78 | 1.11 | 1.03 | 1.25 | 1.06 | 0.89 |
| *pilT* | Twitching motility protein | 2.50 | 1.96 | 2.06 | 1.64 | 2.89 | 1.04 |
| *vgrG-II* | Type VI secretion system protein | 1.55 | 2.41 | 1.78 | 1.33 | 2.23 | 1.84 |
| *ycgB* | Type IV piliation protein | 1.24 | 1.32 | 1.12 | 1.37 | 1.48 | 1.72 |
| *PP_0607* | Type IV pili biogenesis protein FimT | 1.74 | 2.41 | 2.50 | 1.28 | 3.92 | 0.95 |
| *PP_1888* | Fimbrial protein | 1.78 | 1.92 | 2.00 | 1.55 | 1.01 | 1.91 |

| Gene | COG Annotation | Fold Change of FPKM * | | | | | |
|---|---|---|---|---|---|---|---|
| | | Ibuprofen | Naproxen | Gemfibrozil | Diclofenac | Propanolol | Iopromide |
| *PP_4081* | Type IV/VI secretion system protein | 1.17 | 4.32 | 2.60 | 0.93 | 1.74 | 0.96 |

*: Comparing with the control group without pharmaceutical dosage

Table S14. Proteins relevant to fimbriae in recipient bacteria *P. putida* KT2440 after exposure of non-antibiotic pharmaceuticals

| **Protein** | **Description** | **Fold Change of Protein Abundance** * | | | | | |
|---|---|---|---|---|---|---|---|
| | | **Ibuprofen** | **Naproxen** | **Gemfibrozil** | **Diclofenac** | **Propanolol** | **Iopromide** |
| FliC | Flagellin | 1.48 | 1.09 | 0.98 | 1.16 | 1.46 | 1.29 |
| Hcp | Type VI secretion system tube protein | 1.60 | 1.39 | 1.38 | 1.57 | 0.99 | 0.97 |

*: Comparing with the control group without pharmaceutical dosage

Table S15. Genes relevant to SOS response and universal stress in donor bacteria *E. coli* K-12 LE392 after exposure of non-antibiotic pharmaceuticals

| Gene | COG Annotation | Fold Change of FPKM * | | | | | |
|---|---|---|---|---|---|---|---|
| | | **Ibuprofen** | **Naproxen** | **Gemfibrozil** | **Diclofenac** | **Propanolol** | **Iopromide** |
| *lexA* | Transcriptional repressor of SOS regulon | 1.23 | 1.45 | 1.08 | 1.14 | 1.01 | 0.85 |
| *umuC* | Translesion error-prone DNA polymerase V subunit DNA polymerase activity | 1.09 | 1.39 | 5.46 | 1.27 | 1.95 | 0.90 |
| *umuD* | Translesion error-prone DNA polymerase V subunit RecA-activated auto-protease | 3.36 | 2.01 | 1.26 | 1.53 | 3.56 | 1.13 |
| *yebG* | DNA damage-inducible protein regulated by LexA | 1.42 | 1.03 | 0.92 | 1.19 | 1.05 | 0.67 |
| *yebK* | Putative DNA-binding transcriptional regulator | 1.64 | 1.28 | 1.06 | 1.28 | 1.21 | 0.85 |
| *yedK* | DUF159 family protein | 4.96 | 10.34 | 3.39 | 1.21 | 0.55 | 1.04 |
| *uspA* | Universal stress global response regulator | 0.99 | 2.53 | 1.88 | 1.93 | 2.17 | 0.81 |

| **Gene** | **COG Annotation** | **Fold Change of FPKM** * | | | | | |
|---|---|---|---|---|---|---|---|
| | | **Ibuprofen** | **Naproxen** | **Gemfibrozil** | **Diclofenac** | **Propanolol** | **Iopromide** |
| *uspC* | Universal stress protein | 0.67 | 1.56 | 1.62 | 1.12 | 1.27 | 0.64 |
| *uspD* | Stress-induced protein | 0.55 | 2.01 | 1.44 | 1.36 | 1.59 | 1.05 |
| *uspE* | Stress-induced protein | 1.12 | 2.10 | 1.61 | 1.48 | 1.84 | 0.94 |
| *uspF* | Stress-induced protein ATP-binding protein | 1.16 | 1.25 | 1.27 | 0.72 | 1.09 | 1.21 |
| *uspG* | Universal stress protein UP12 | 0.77 | 1.77 | 1.55 | 1.21 | 1.53 | 1.09 |

*: Comparing with the control group without pharmaceutical dosage

Table S16. Proteins relevant to SOS response and universal stress in donor bacteria *E. coli* K-12 LE392 after exposure of non-antibiotic pharmaceuticals

| **Protein** | **Description** | **Fold Change of Protein Abundance** * | | | | | |
|---|---|---|---|---|---|---|---|
| | | **Ibuprofen** | **Naproxen** | **Gemfibrozil** | **Diclofenac** | **Propanolol** | **Iopromide** |
| UspG | Universal stress protein UP12 | 1.48 | 1.93 | 1.61 | 1.36 | 1.21 | 0.84 |

*: Comparing with the control group without pharmaceutical dosage

Table S17. Genes relevant to SOS response and universal stress in recipient bacteria *P. putida* KT2440 after exposure of non-antibiotic pharmaceuticals

| **Gene** | **COG Annotation** | **Fold Change of FPKM** * | | | | | |
|---|---|---|---|---|---|---|---|
| | | **Ibuprofen** | **Naproxen** | **Gemfibrozil** | **Diclofenac** | **Propanolol** | **Iopromide** |
| *soxD* | Sarcosine oxidase subunit delta | 1.60 | 2.55 | 1.24 | 0.52 | 1.64 | 1.55 |
| *soxR* | DNA-binding transcriptional regulator | 2.00 | 1.69 | 1.69 | 3.94 | 4.17 | 4.08 |
| *PP_2326* | Universal stress protein | 1.19 | 1.43 | 1.91 | 1.26 | 1.57 | 0.91 |
| *PP_3288* | Universal stress protein family protein | 1.23 | 1.57 | 1.55 | 1.33 | 1.47 | 1.83 |

*: Comparing with the control group without pharmaceutical dosage

Table S18. Proteins relevant to SOS response and universal stress in recipient bacteria *P. putida* KT2440 after exposure of non-antibiotic pharmaceuticals

| **Protein** | **Description** | **Fold Change of Protein Abundance** [*] | | | | | |
|---|---|---|---|---|---|---|---|
| | | **Ibuprofen** | **Naproxen** | **Gemfibrozil** | **Diclofenac** | **Propanolol** | **Iopromide** |
| NP_745431.1 | Universal stress protein family protein | 1.18 | 1.08 | 1.01 | 1.56 | 1.01 | 1.04 |

[*]: Comparing with the control group without pharmaceutical dosage

Table S19. Genes relevant to efflux pump and repressor to antibiotic sensitivity in donor bacteria *E. coli* K-12 LE392 after exposure of non-antibiotic pharmaceuticals

| **Gene** | **COG Annotation** | **Fold Change of FPKM** * | | | | | |
|---|---|---|---|---|---|---|---|
| | | **Ibuprofen** | **Naproxen** | **Gemfibrozil** | **Diclofenac** | **Propanolol** | **Iopromide** |
| *aaeX* | DUF1656 family putative inner membrane efflux pump associated protein | 4.66 | 1.16 | 1.48 | 2.13 | 1.26 | 2.31 |
| *mdtJ* | Multidrug efflux system transporter | 1.85 | 1.27 | 1.35 | 3.71 | 2.31 | 2.95 |
| *yhiI* | Putative membrane fusion protein (MFP) of efflux pump | 1.21 | 1.06 | 3.68 | 1.57 | 1.75 | 2.22 |
| *kdgR* | KDG regulon transcriptional repressor | 2.00 | 0.99 | 1.39 | 1.20 | 1.23 | 1.11 |

*: Comparing with the control group without pharmaceutical dosage

Table S20. Genes relevant to efflux pump in recipient bacteria *P. putida* KT2440 after exposure of non-antibiotic pharmaceuticals

| Gene | COG Annotation | Fold Change of FPKM * | | | | | |
|---|---|---|---|---|---|---|---|
| | | **Ibuprofen** | **Naproxen** | **Gemfibrozil** | **Diclofenac** | **Propanolol** | **Iopromide** |
| *czcA-I* | Cation efflux system protein | 1.68 | 2.77 | 1.23 | 1.37 | 1.37 | 1.68 |
| *czcA-II* | Cation efflux system protein | 2.08 | 1.91 | 1.22 | 1.01 | 0.81 | 0.78 |
| *PP_3789* | Efflux transporter | 1.16 | 2.73 | 2.17 | 1.26 | 4.53 | 1.29 |
| *PP_0805* | Outer membrane efflux protein | 1.09 | 1.16 | 2.19 | 1.31 | 1.89 | 0.85 |
| *PP_1152* | Membrane fusion efflux protein | 1.21 | 3.03 | 1.07 | 1.39 | 1.41 | 0.61 |
| *PP_4923* | Outer membrane efflux protein | 1.47 | 0.90 | 1.44 | 1.45 | 1.23 | 0.85 |

*: Comparing with the control group without pharmaceutical dosage

Table S21. Primers used in this study [1,5]

| Gene | Primer | Sequence of primer |
|---|---|---|
| *tetA* | Short FW | GACTATCGTCGCCGCACTTA |
| | Short RV | ATAATGGCCTGCTTCTCGCC |
| | Long FW | CGTGTATGAAATCTAACAATGCGCT |
| | Long RV | CCATTCAGGTCGAGGTGGC |
| *bla* | Short FW | AATAAACCAGCCAGCCGGAA |
| | Short RV | TTGATCGTTGGGAACCGGAG |
| | Long FW | TTACCAATGCTTAATCAGTGAGGC |
| | Long RV | ATGAGTATTCAACATTTCCGTGTCG |
| *traF* | FW | AAGTGTTCAGGGTGCTTCTGC |
| | RV | GTCGCCTTAACCGTGGTGTT |
